# Mathematically mapping the network of cells in the tumor microenvironment

**DOI:** 10.1101/2023.02.03.526946

**Authors:** Mike van Santvoort, Óscar Lapuente-Santana, Francesca Finotello, Pim van der Hoorn, Federica Eduati

**Affiliations:** Department of Mathematics and Computer Science, Eindhoven University of Technology, Eindhoven, PO Box 513, 5600MB, Eindhoven, The Netherlands; Institute for Complex Molecular Systems, Eindhoven University of Technology, PO Box 513, 5600MB, Eindhoven, The Netherlands; Department of Biomedical Engineering, Eindhoven University of Technology, PO Box 513, 5600MB, Eindhoven, The Netherlands; Universität Innsbruck, Department of Molecular Biology, Digital Science Center (DiSC), Innrain 52, 6020 Innsbruck, Austria

## Abstract

Cell-cell interaction networks are pivotal in cancer development and treatment response. These networks can be inferred from data; however, this process often combines data from multiple patients, and/or creates networks on a cell-types level. It creates a good average overview of cell-cell interaction networks but fails to capture patient heterogeneity and/or masks potentially relevant local network structures. We propose a mathematical model based on random graphs (called RaCInG) to alleviate these issues using prior knowledge on potential cellular interactions and patient’s bulk RNA-seq data. We have applied RaCInG to extract 444 network features related to the tumor microenvironment, unveiled associations with immune response and subtypes, and identified cancer-type specific differences in inter-cellular signaling. Additionally, we have used RaCInG to explain how immune phenotypes regulated by context-specific intercellular communication affect immunotherapy response. RaCInG is a modular pipeline, and we envision its application for cell-cell interaction reconstruction in different contexts.

## Introduction

In the fight against cancer, it is key to stratify patients based on tumor characteristics, since these predict how a patient will respond to treatment. To stratify effectively, one needs to measure the functional state of the cells and molecules that reside in a tumor, collectively called the tumor microenvironment (TME). Big breakthroughs have been achieved focusing on the functionality of individual cells and proteins. For example, the development of programmed cell death ligand 1 (PD-L1) blockers^1^ to counteract the protein’s unambiguous pro-tumor effect^2^.

However, the TME exhibits emergent behavior that cannot be explained by individual cell- or protein types^3,4^ and focusing only on individual parts of the TME hinders the development of more comprehensive treatment strategies. For example, the tumor necrosis factor alpha (TNF-α) protein can elicit both a pro- or anti-tumor reaction based on further context cues in the TME^5^. Thus, to fully capture the functional state of the TME it should be considered as an interconnected system rather than a collection of individual components.

An unbiased approach to do this consists in the modeling of the TME as a cell-cell communication network, which can be inferred typically from RNA sequencing (RNAseq) data using statistical inference methods or machine learning techniques^6^. Several studies have shown the value of using the reconstructed cell-cell communication networks to study the role of cell-cell communication in the TME^7–12^. However, existing techniques have several drawbacks. Most of them build a network on the cell- and protein-type level and not on the level of individual cells/proteins^6–9^. This creates a “low resolution picture” of the cell-cell communication network that masks important local network structures. Moreover, these methods are often do not capture cell-cell communication networks of individual patients^11,13^.

Most of the methods that construct networks of individual cells or individual patients rely on single-cell RNA-sequencing (scRNA-seq) data to derive their networks^9,12^. This provides “higher resolution” modeling, but is more complicated to apply in specific use cases, since scRNA-seq data itself has some technical limitations: higher uncertainty, drop-outs, and limited clinical applicability due to its higher costs and difficulties in sample preparation^14^. A recent approach has been proposed that combines bulk RNA-seq data with probabilistic techniques to reconstruct cell-cell interaction networks for individual patients^10^. However, this method builds a network on the level of cell-types that provides only a mean-field approximation to the actual cell-cell interaction network without mathematical guarantees on how well this approximation fits the data.

The field of random graphs models^15^ can help in addressing these limitations, providing natural ways to deal with limited prior knowledge when constructing cell-cell interaction networks. Where prior knowledge fails us, stochasticity of random graphs can fill the knowledge gaps in the most unbiased way possible, ensuring the result has no statistical bias outside the provided data. Although these models explicitly introduce noise in the cell-cell interaction network construction, emergent network behavior remains statistically consistent. These consistencies can be mathematically proven, extracted and used as fingerprints of the actual cell-cell interaction network. We can then use these fingerprints as features to understand, predict and ultimately reshape the TME. Thus, even if we cannot derive a cell-cell communication network at the level of individual cells directly from available data, random graph modeling will still allow us to pinpoint local properties that should emerge from such “high resolution” networks. In practice, this means random graph modeling allows us to reconstruct single cell networks from widely accessible bulk RNA-seq data. It can do this by relying on prior knowledge until uncertainties are encountered, which it resolves by sampling from all possible options uniformly at random without making extra explicit assumptions^16^.

Here, we provide a methodological pipeline to reconstruct (ensembles of) cell-cell interaction networks using patient-specific bulk RNA-seq data and prior knowledge on the ligands and receptors that can be secreted by given cell-types as input. We make use of a specifically designed random graph model as a statistical model for the potential configurations of the network that respects constraints from the input data and biology. We provide this data analysis pipeline as a general toolbox called the “Random Cell-cell Interaction Generator” (RaCInG) to study context-specific cell-cell interaction networks. To validate the pipeline, we reconstruct cell-cell interactions among relevant cell-types in the TME for 3310 cancer patients. In these case studies we show that we can extract consistent properties from individual patients that form predictors for their immune subtype and response to immunotherapy with immune checkpoint blockers (ICB).

## Results

### Reconstructing cell-cell communication networks through monte-carlo simulations

RaCInG constructs directed networks where the nodes represent individual cells, and the arcs (i.e., the directed connections) represent ligand-receptor interactions between two cells. To generate networks, four types of input are needed (**Fig. 1A**): 1. A cell-type vector (*C-distribution*), where each entry indicates the probability of an individual cell having a given type. 2. A ligand by receptor matrix (*LR-distribution*), where each entry indicates the probability of an individual interaction involving a given ligand and receptor. 3 and 4. A ligand (or receptor) by cell-type binary matrix, where 0 indicates that a ligand (or receptor) cannot be expressed by a cell-type, and 1 indicates that it can. An example of how these input matrices can be derived from patient-specific bulk RNA-seq data and general prior knowledge is provided later in the case study.

**Fig. 1:**
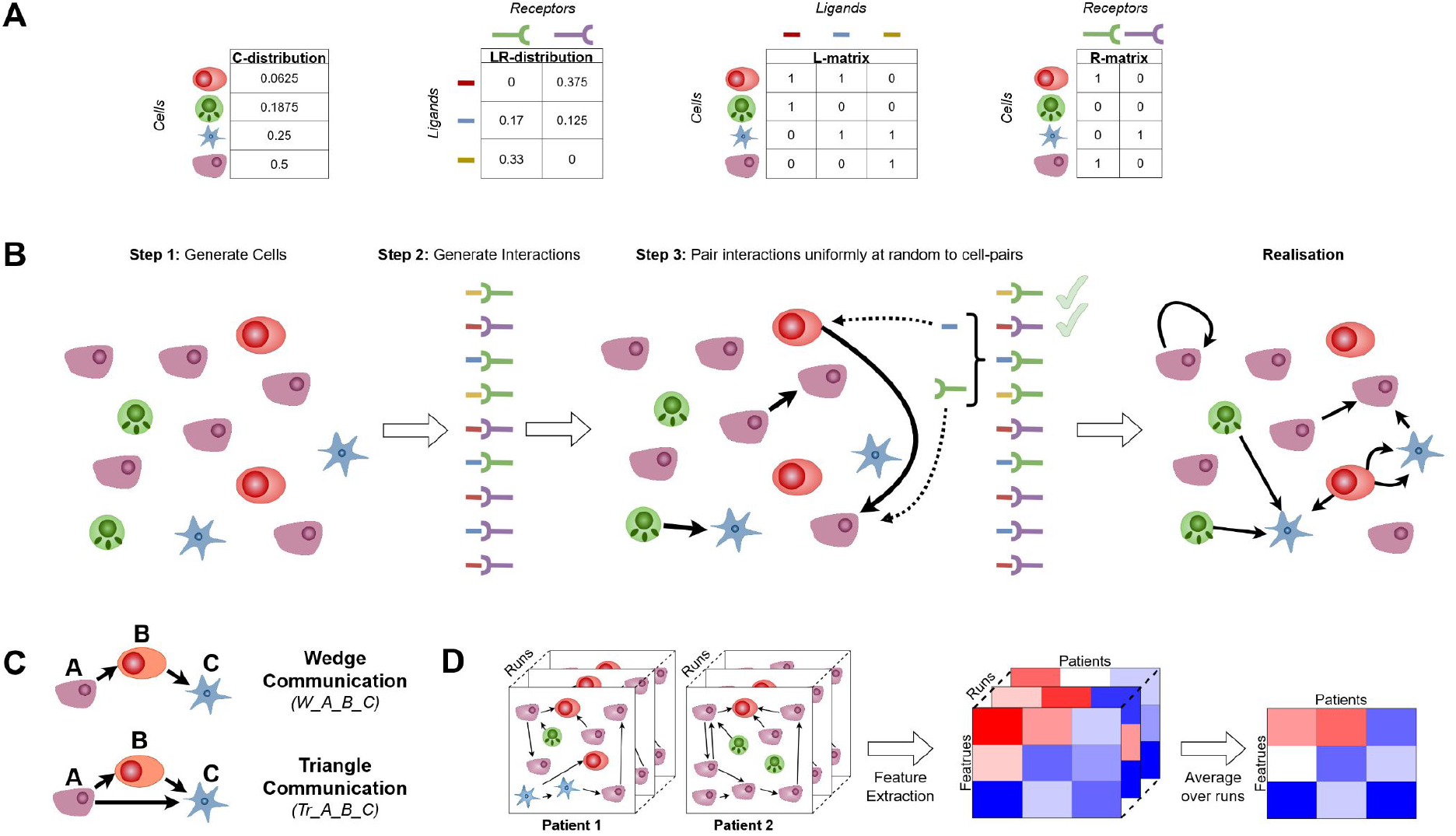
Methodology of monte-carlo simulation. (A) Input matrices used by RaCInG including information on: cell-types and ligand-receptors relative quantification (*C-distribution* and *LR-distribution* respectively); which ligands and receptors can be expressed by specific cell types (L-matrix and R-matrix respectively). (B) Schematic depiction of the simulation steps for one network based on the input matrices including: random generation of cells and ligand-receptor interactions based on *C-distribution* and *LR-distribution* matrices respectively (Step 1-2); iterative assignment of ligand-receptor interactions to cell-pairs based on *L-matrix* and *R-matrix* (Step 3). (C) The types of features extracted from the simulated networks. (D) The global pipeline of the monte-carlo method including the generation of multiple possible realizations of random networks and the extraction of robust network features.

The network generation procedure (**Fig. 1B**) starts by generating a fixed number of individual cells whose types are assigned randomly based on the *C-distribution*. Then, a fixed number of random ligand-receptor interactions are generated based on the *LR-distribution*. Treating the C- and LR-distributions as probabilities rather than as exact numbers allows handling uncertainties in the input data.

The procedure continues by attaching each ligand-receptor interaction as an arc in between two cells selected uniformly at random among the ones which can express the ligand and the receptor as defined through the *L-matrix* and *R-matrix*, respectively. This process of connecting cell-cell pairs based on ligand and receptors continues until all ligand-receptor pairs have been assigned.

After this procedure, RaCInG has created one network instance for a given patient that adheres to the constraints from RNA-seq data. This is only one possible representation of the network and is not necessarily representative of the patient’s actual cell-cell communication network. Thus, RaCInG generates an ensemble of networks for the same patient and extracts statistical properties that remain consistent in the network ensemble^17^. We define these as network fingerprints which include information about high-level interactions between two or more cell-types (graphlets) and about low-level interactions between ligands and receptors.

Currently, two types of fingerprints involving triplets of cells are extracted based on monte-carlo simulations by RaCInG: wedges and triangles (**Fig. 1C**). Specific wedges and triangles are referred to hereafter as *W_A_B_C* and *Tr_A_B_C*, respectively, with letters indicating the cell types involved. The count of wedges and triangles for individual patient-specific networks is computed as the average over the ensemble to account for model randomness and derive close approximation of their abundance in the actual cell-cell communication network (**Fig. 1D**, see **Methods** for the computation of the counts). The quantification of network fingerprints for individual patients are interpreted as features for further analysis.

### Kernel-based approach to derive network fingerprints

Although monte-carlo simulation provides an intuitive method to extract features, RaCInG also allows to mathematically derive some features using random graph theory, based on kernels. This is a matrix that encodes the asymptotic probability that a ligand-receptor interaction exists between two individual cells with specific cell-types. It is based on the expected number of ligand-receptor interactions that connect these cells^15,18–20^ (see **Methods** for the exact expression). If we would generate networks using a kernel, then after generation of cells a coin flip determines whether a ligand-receptor interaction between each pair of individual cells appears. The success probability of this flip is determined by the cell-types of the pair and their kernel value (**Fig. 2A**).

**Fig. 2:**
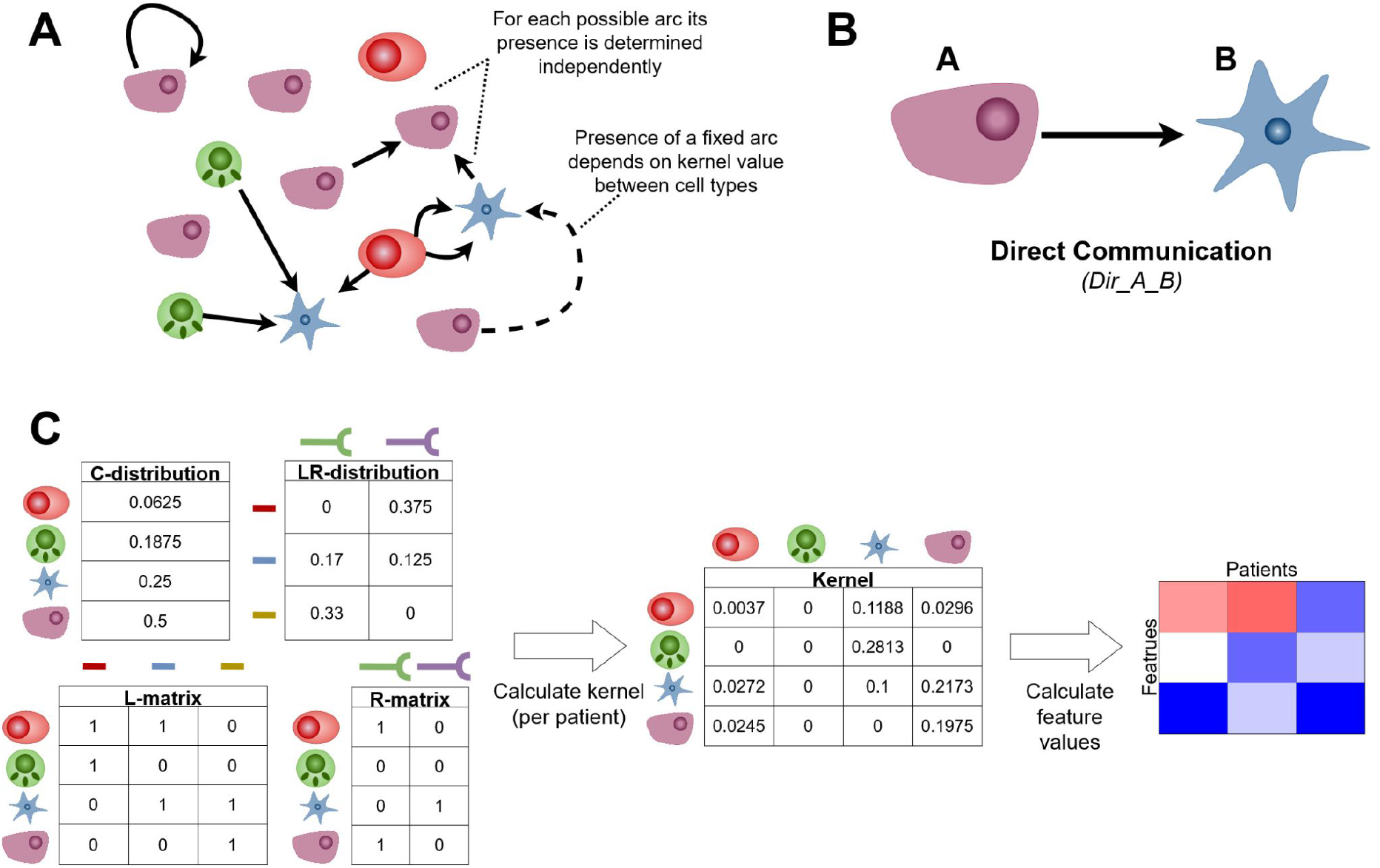
Kernel method based on random graph theory. (A) The mathematical idea behind graph generation in the kernel method, which highlights the interpretation of the kernel. (B) The feature type extracted using the kernel method. (C) The general pipeline used to extract features from the graph using the kernel method.

RaCInG allows quantifying the direct communication between individual cells with cell-type A and B (referred hereafter as *Dir_A_B*, **Fig. 2B**) using the kernel method. First the kernel is computed for each patient using all four input matrices, and then the kernels are transformed into the feature values (**Fig. 2C**). This approach uses a mathematical guarantee (see **Methods** for the derivation) and is faster than using the network generation procedure.

### Normalization of network fingerprints to account for different cellular composition

All methodologies in RaCInG to extract network features (i.e., the monte-carlo and kernel method) are biased through cell-type quantification. When assigning ligand-receptor pairs to cells, the model selects cells uniformly at random, so highly abundant cells-types have a larger probability of being selected, reflecting in the feature values. To account for this and allow comparison of network features between samples with different cellular composition, we implement a normalization procedure that corrects for the influence of the cell-type abundance.

RaCInG recomputes the network features for each patient using the same input matrices except for the *LR-distribution*, which is made uniform (i.e., same probability for all ligand-receptor interactions). This removes the influence of the ligand-receptor quantification, as the features extracted using the uniform *LR-distribution* are determined just by cell-type quantification. Finally, we compute the fold-change between the (average) feature values obtained using the data-derived versus uniform *LR-distributions*. The resulting feature values depend predominantly on ligand-receptor quantification. Moreover, the procedure ensures that all feature values have the same order of magnitude, regardless of their type.

### Application to characterize the tumor microenvironment

We used RaCInG to investigate the role of cell-cell communication by building patient-specific cell-cell interaction network models for 3213 patients from six solid cancers from The Cancer Genome Atlas (TCGA): bladder urothelial carcinoma (BLCA; N = 407), colon rectal cancer (CRC; N = 379), clear cell renal cell carcinoma (KIRC; N = 533), non-small cell lung cancer (NSCLC; N = 1012), skin cutaneous melanoma (SKCM; N = 467) and stomach adenocarcinoma (STAD; N = 415) (**Methods**)^21^.

We first derived the four input matrices required by RaCInG as follows (**Fig. 3A**; see **Methods** for more details): 1. The *C-distribution* table consists of nine cell-types present in the TME (names and abbreviations are summarized in **Table 1**). The probability of each cell-type to appear in the network was defined specifically for each patient from RNA-seq data as their relative abundance quantified using an ensemble of deconvolution methods^22^. The *LR-distribution* table was defined based on a list of 971 literature-curated LR interactions^23^ and quantified for each patient as the most limiting factor between the expression of the ligand and the receptor based on the RNA-seq data. 3-4. The *L-matrix* and the *R-matrix* were defined as (non patient-specific) prior knowledge that indicated which ligand and receptor can be expressed by a specific cell-type based on cell-type specific gene expression data^24^.

**Fig. 3:**
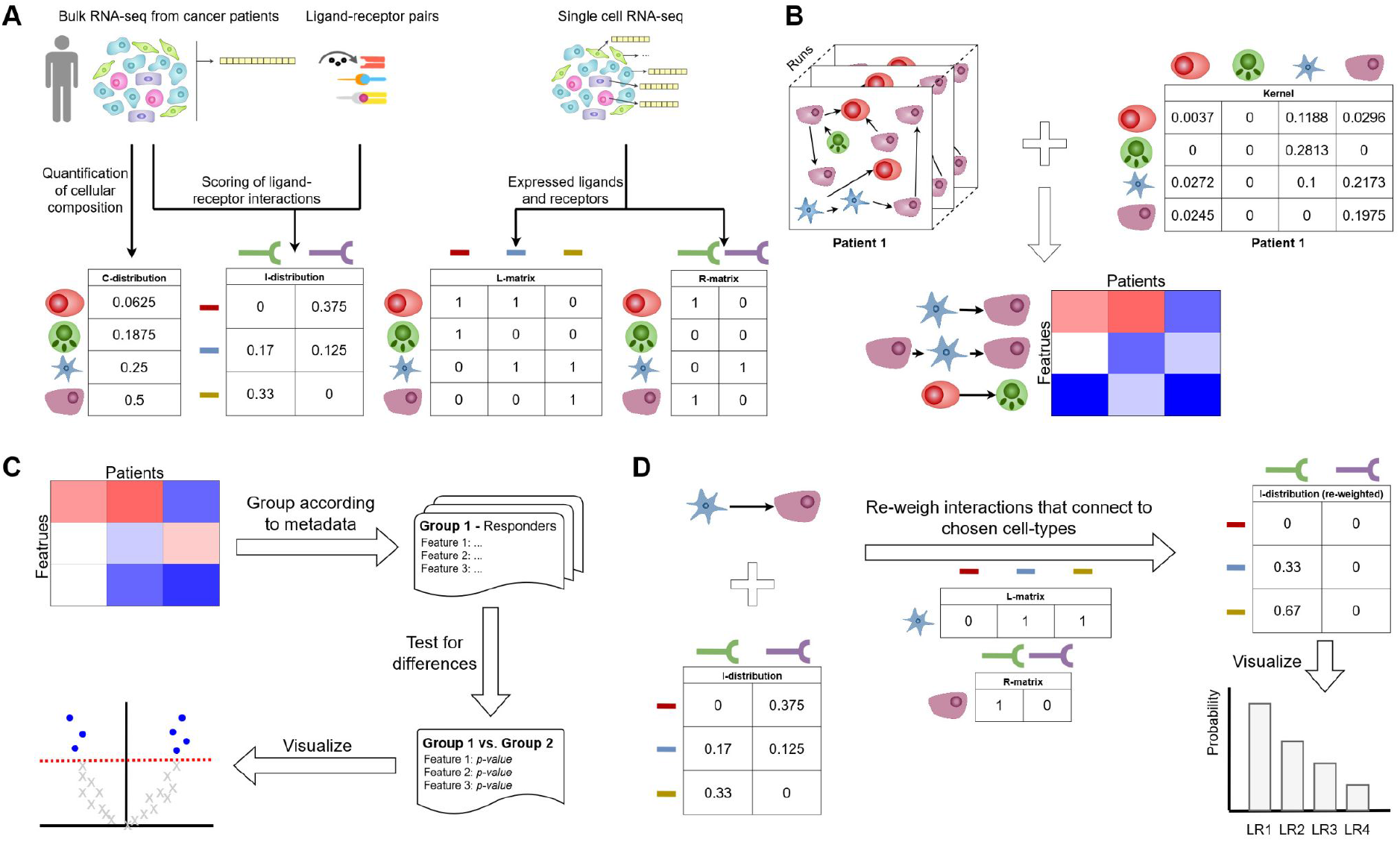
Modular structure of RaCInG for analysis of cell-cell interactions in the TME. (A) Cell- and interaction-quantification from bulk RNA-seq data. (B) Feature extraction from patient specific graphs or kernel values. (C) Statistical analysis based on a list of extracted feature values in a batch of patients. (D) Extraction of LR-pair probabilities for given cell-type interactions.

**Table 1:** The cell-types included in our case studies.

| Cell name | Abbreviation | Cell name | Abbreviation |
| --- | --- | --- | --- |
| Tumor | *Tumor* | B-cell | *B* |
| Cancer associated fibroblast | *CAF* | Macrophage | *M* |
| Endothelial cell | *Endo* | Dendritic cell | *DC* |
| CD8+ T-cell | *CD8* | Regulatory T-cell | *Treg* |
| Natural killer cell | *NK* |  |  |

Using the monte-carlo simulations and the kernel method described above (**Fig. 1D** and **2C**) we derived the three sets of network features consisting of: 81 direct communications, 729 wedge communications and 978 triangle communications (**Fig. 3B**). Based on our first analysis of the results, showing a limited influence of the directionality of interactions, we decided to consider classes of undirected interactions (**Methods**). Additionally, due to the low and inconsistent quantification of NK cells, features involving this cell-type were discarded. These adjustments reduced the number of network features to 36, 288 and 120, respectively. Finally, we used the extracted features (**Supplementary Table 1**) to compare patients or patient groups (**Fig. 3C**) and look into the LR-pairs that make up specific features of interest (**Fig. 3D**).

### Network features correlate with immune response

Cell-cell communication has shown to influence the orchestration of anti-cancer immune response^25^. Therefore, we have applied RaCInG to investigate how different graph features in our models correlate with an ensemble immune response score (**Methods**)^26^.

We observed that 31-87 features out 444 (7-20% depending on cancer type) strongly correlated with immune response (absolute Spearman rho > 0.5; p-val < 0.01 after Bonferroni correction; **Fig. 4**). Generally, the more “complex” features (i.e., wedges and triangles) showed similar associations with immune response (6.6%–18.1% and 6.7%-21.7% highly correlated features respectively) as the “simple” fingerprints (i.e., direct communication; 8.3%-38.9% highly correlated features). However, if a direct communication feature appeared (e.g. *Dir_CD8_M* in NSCLC; rho = 0.652, p-val < 0.0001), then often a more complex feature, including this direct communication as a subset, showed a higher absolute correlation (e.g. *W_CD8_M_M;* rho = 0.754, p-val < 0.0001). Such more complex features are more informative as they describe intercellular communication paths rather than simple direct interactions and they can highlight which detailed interaction elicits the strong correlation with immune response. Following the example above, *W_CAF_M_CD8* has lower correlation (rho = 0.590, p-val < 0.0001) meaning that the addition of CAFs to direct communication between macrophages and T-cells worsens the overall immune response. These observations highlight the importance of looking at more intricate communication mechanisms to study the coordination of anti-cancer immune responses.

**Fig. 4:**
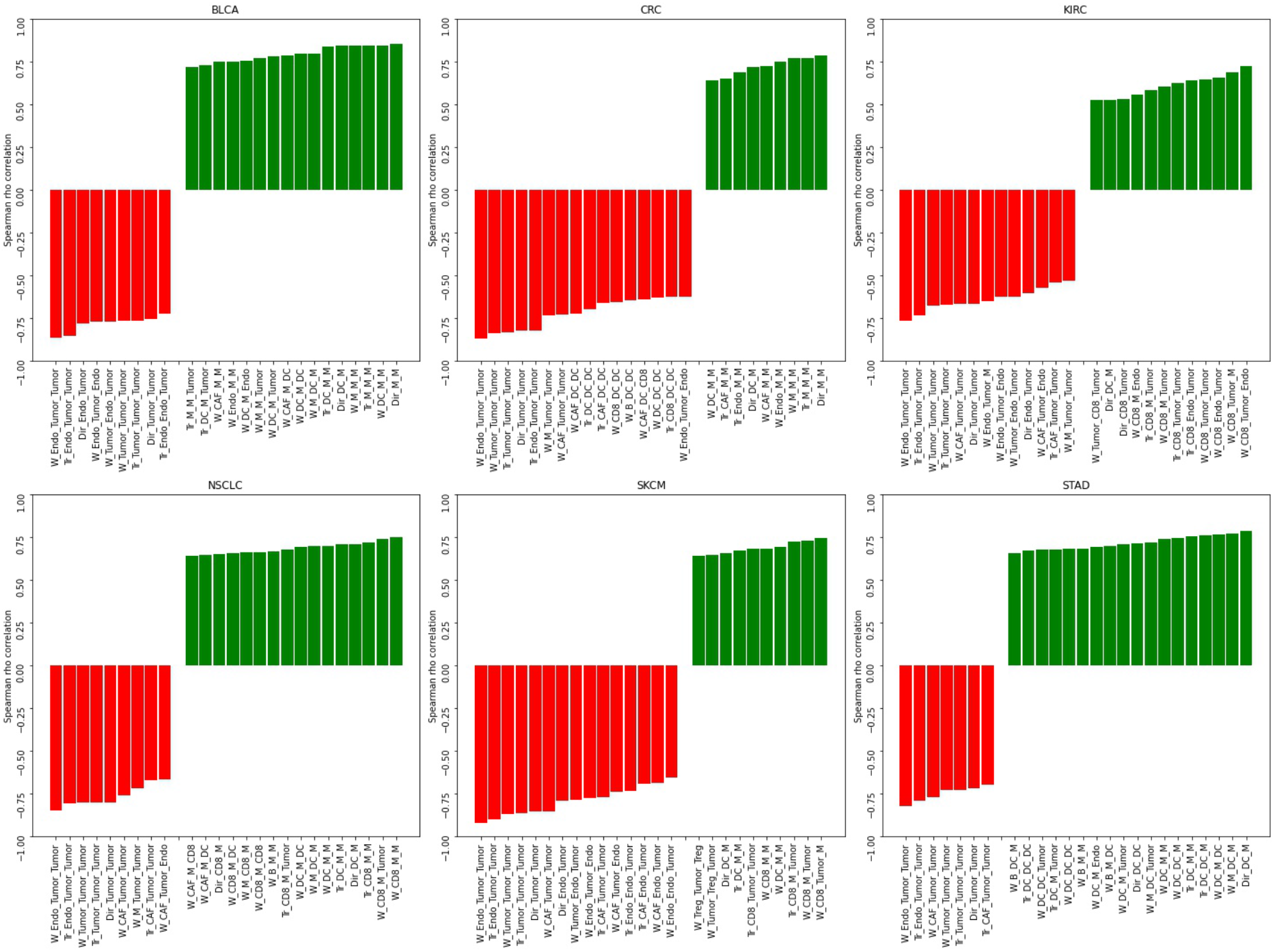
Network features associated with immune response. The 25 features with the largest spearman rho correlation with immune response for each of the six cancer types. Features were only selected if their associated p-value was smaller than 0.01 after Bonferroni correction.

When focusing on features that showed negative association with immune response, we mainly found communication structures consisting of tumor cells, endothelial cells and CAFs and absence of involvement of immune cells. One reason for that is the direct positive relationship between immune response and the presence of infiltrated immune cells in the tumor (e.g., through the formation of tertiary lymphoid structures)^27,28^. In this scenario, we expect the immune cells to drive the communication with the aforementioned three cell-types. Instead, in tumors with a less active immune response, the main remaining communication players will be tumor cells, CAFs and endothelial cells.

Interestingly, we observed a different behavior in CRC, where endothelial cell and tumor communication as well as dendritic cell communication to CAFs and CD8+ T-cells were negatively correlated with the immune response score (**Fig. 4**; CRC panel). This can be explained by the fact that dendritic cells in CRC are mostly found in the tumor stroma and are therefore likely to establish crosstalks with stromal cells like CAFs^29^. In agreement with the negative correlation, dendritic cells in the stroma have been shown to be associated with low infiltration of CD8+ T-cells^29^.

Concerning positive associations with immune response, we observed a varied palette of features, with communication predominantly between immune cells or between immune cells and tumor cells found among the features with the largest, positive correlation. The specific set of features correlated with immune response varies between cancer types, confirming the well-established heterogeneity of the TME across cancer types^30^. This heterogeneity underlies the potential of deriving patient-specific models of intercellular communication, as we will further explore in the next sections.

### Network features as immune phenotype indicators

To uncover the existent heterogeneity of cell-cell communication across patients, we used RaCInG to seek whether certain network features can explain differences between immune phenotypes. We considered four immune phenotypes previously defined in literature^31^: immune enriched (IE), immune enriched-fibrotic (IE/F), fibrotic (F) and immune deprived (D). The immune enriched groups (IE and IE/F) are characterized by high anti-tumor immune cell infiltration and activation in the tumor. The fibrotic groups (IE/F and F) are characterized by activation of stromal cells like CAFs. IE and F tumors are expected to have positive and negative correlation with response to ICB therapy respectively. Finally, the deprived group (D) is characterized by little immune or stromal cell activation. In the following sections we start from a pan-cancer analysis of patients from the six cancer types discussed above, and then focus on two cancer-specific analyses.

#### Pan-cancer analysis

When looking at comparison between immune subtypes at the pan-cancer level (**Fig. 5**) we identified that the D phenotype is mainly characterized by communication between tumor and endothelial cells. This is in agreement with the expected negative association with the immune response. The D group is likely to have minimal leukocyte or lymphocyte activity^31^, opening the door for high cellular communication between malignant and non-immune cell-types.

**Fig. 5:**
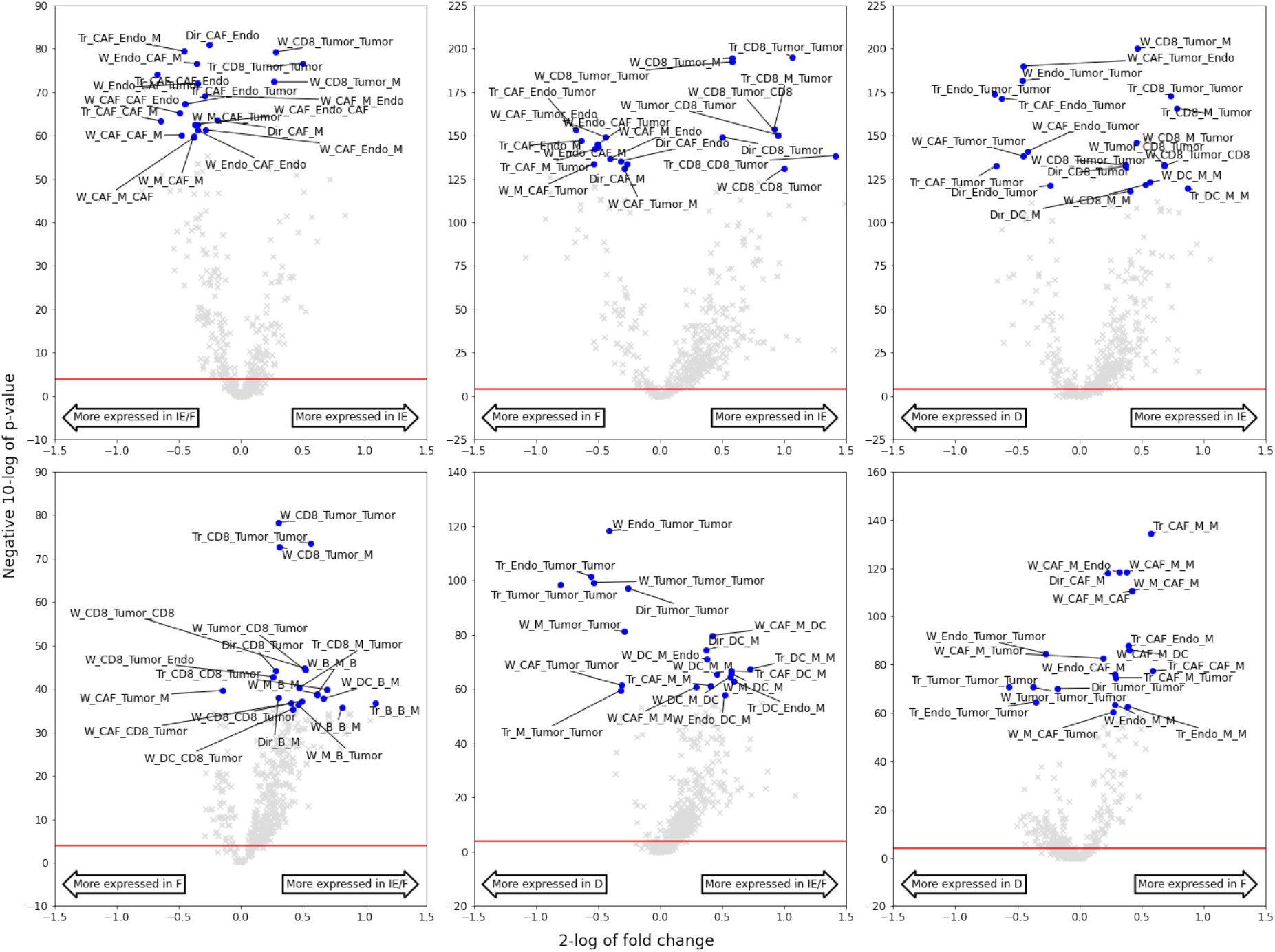
Network-based characterization of microenvironment subtypes in pan-cancer settings. Volcano plots showing the statistical comparison of network-based features identified by RaCInG when doing pairwise comparisons of microenvironment subtypes across cancer types. The red line indicates the α = 0.05 significance threshold after Bonferroni correction. On the x-axis we show the fold change between the average feature values for each group, and on the y-axis the negative 10-log of the Wilcoxon rank sum test’s p-value. For each plot, the twenty features with lowest p-value have been highlighted.

As expected, we identified increased CAF activity in the fibrotic groups (IE/F and F) as well as CD8+ T-cell activity in the immune enriched groups (IE and IE/F). Interestingly, we see in the IE to F comparison that macrophages appear in both groups: communicating with CD8+ T-cells in the IE group, or with CAFs in the F group. The dual importance of macrophages in both IE and F subtypes might be explained by macrophages playing different roles in the tumor depending on their phenotype. Anti-tumor macrophages (also called M1 macrophages) recruits CD8+ T-cells to fight the tumor^32^, explaining its appearance in the IE group where we expect to have higher anti-tumor to pro-tumor macrophage ratio^31^. On the contrary, pro-tumor macrophages (also called M2 macrophages) are known to conspire with CAFs to boost tumor malignancy^33,34^, motivating why this interaction appears in the most hostile immune phenotype. Overall, these considerations show that the network features are able to capture general characteristics of immune phenotypes well and show potential to distinguish the functional role of cell-types based on their interactions.

#### Cancer type-specific analysis

Next, we focused our analysis on melanoma (SKCM) and gastric cancer (STAD) to show which additional insights RaCInG can provide at the cancer-specific level (**Fig. 6**; **Supplementary Fig. 1-3**).

**Fig. 6:**
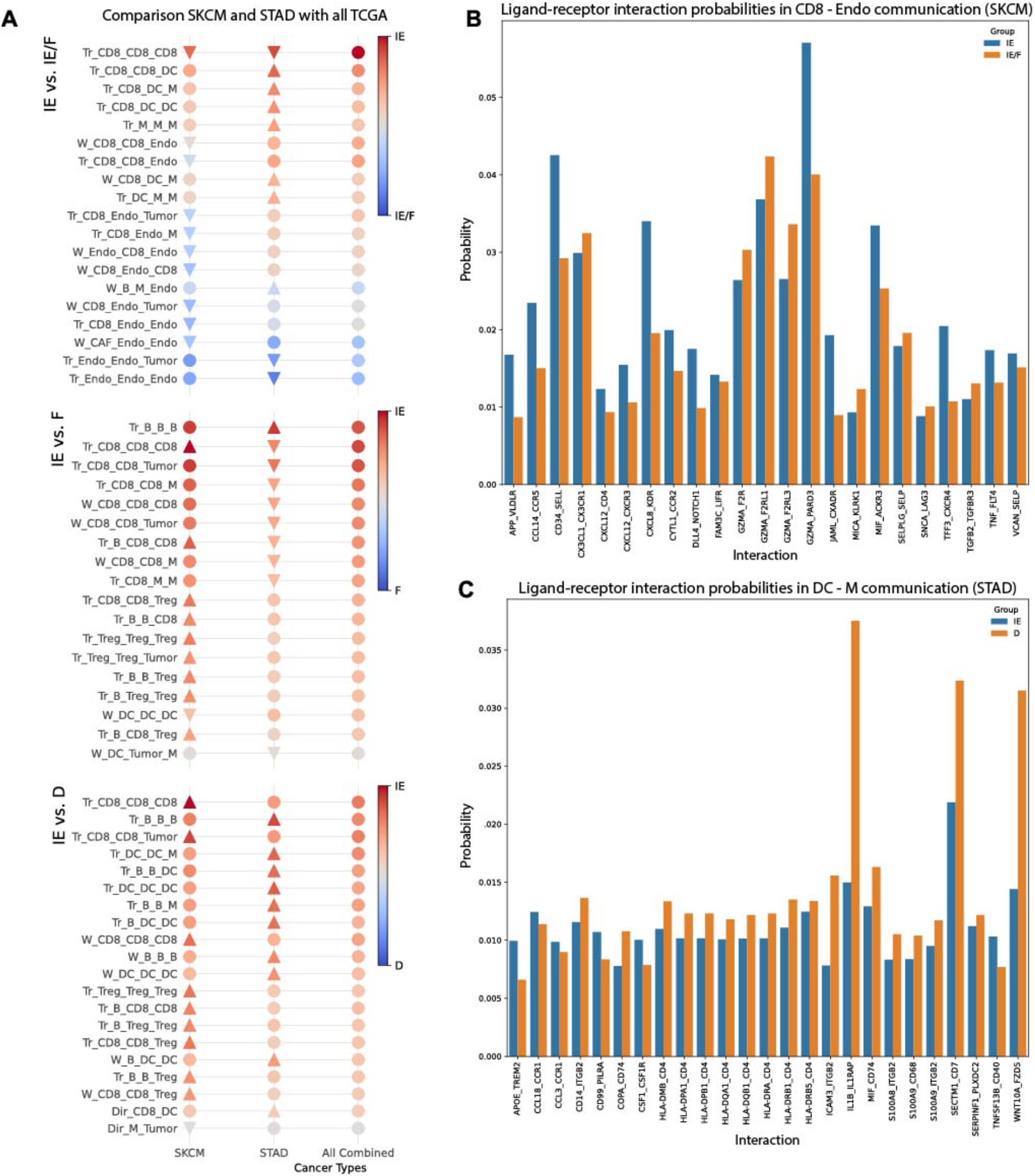
Cancer specific analysis of SKCM and STAD. (A) List of top ten features for SKCM and STAD that changed the most in average fold-change when compared to the results of the pan-cancer analysis. The direction of the triangles indicates the direction of the fold-change shift when compared to the pan-cancer analysis. Dots indicate that the fingerprint is not part of the top ten features for the given cancer type. (B) The top twenty ligand-receptor pairs that are most likely to create a connection between CD8+ T-cells and endothelial cells in SKCM for the immune subtypes IE and IE/F. (C) The top twenty ligand-receptor pairs that are most likely to create a connection between dendritic cells and macrophages in STAD for the immune subtypes IE and D.

When comparing the IE versus non-IE (F and D) subtypes we observed that several features associated with immune activation are more prominent in SKCM compared to pan-cancer (**Fig. 6A**). Examples are B-cells activating CD8+ T-cells (e.g. Tr_B_CD8_CD8, 2-log fold-change 0.99 and 1.34 in IE vs D and IE vs F comparisons respectively), self-activation of CD8+ T-cells (e.g. Tr_CD8_CD8_CD8, fold-change 1.66 and 1.77 in IE vs D and IE vs F comparisons respectively) and CD8+ T-cells targeting tumor cells (*Tr_CD8_CD8_Tumor*, fold-change 1.40 in the IE vs D comparison). These observations are in agreement with the strong immune response reported in SKCM^31^. This strong antitumor immune response can cause the recruitment of immunosuppressive T_re_g cells by CD8+ T-cells (*Tr_CD8_CD8_Treg;* fold-change 1.08 and 1.14 in IE vs D and IE vs F comparisons respectively) and B-cells (*Tr_B_B_Treg*; fold-change 0.90 and 1.03 in IE vs D and IE vs F comparisons respectively) to counterbalance the high immunogenicity of these tumors and as a potential mechanism of immune evasion^35–37^.

When comparing IE and IE/F groups we observed that the CD8+ T-cell communication with endothelial cells is stronger in IE/F patients for the SKCM dataset when compared to the pan-cancer analysis (eight out of the top ten features that are more specific for SKCM involve this interaction, all with 2-log fold-change < 0; arrows pointing down in **Fig. 6A** for SKCM). Often, these features shifted from being more represented in IE patients to being enriched in IE/F patients (see e.g., *W_CD8_Endo_CD8;* going from red in the pan-cancer to blue in the SKCM comparison of IE vs IE/F in **Fig. 6A**).

To delve deeper into what proteins contribute to this shift, we retrieved from RaCInG the top 20 ligand-receptor interactions that are likely to drive this cell-cell communication (**Fig. 6B**). Three out of the five interactions with a higher probability of appearing in the IE/F subtype compared to the IE subtype involve a member of the family of thrombin receptors (F2R, F2RL1, and F2RL3) interacting with granzyme A (GZMA). Interestingly, GZMA interacting with thrombin receptors is usually associated with apoptosis in targeted cells^38,39^, creating an anti-tumor microenvironment that is more fitting for the IE subtype. However, in melanoma thrombin receptors stimulation has been associated with tumor progression, which is more common in IE/F patients^40^.

Regarding STAD, we observed that interactions involving dendritic cells, especially with macrophages, are the most distinguishing features which are downregulated in the D subtype compared to pan-cancer (arrows going up in the IE vs D comparison **Fig. 6A**). Focusing on the interaction between macrophages and dendritic cells, we identified three ligand-receptor pairs which are particularly more abundant in the D than in the IE subtype in STAD (**Fig. 6C**).

These are the interactions between interleukin 1 beta (IL1B) and interleukin-1 receptor accessory protein (IL1RAP), between WNT family member 10A (WNT10A) and frizzled class receptor 5 (FZD5), and between secreted and transmembrane protein 1 (SECTM1) and CD7. By communicating through the IL1RAP and IL1B proteins, the macrophages and dendritic cells dampen the inflammatory process in the D subtype (if they communicate), inducing a poor prognosis^41^. This entails that in the D subtype immune cells interact less, explaining why globally we see macrophage interaction with dendritic cells more in the IE subtype, where inflammation is stronger. Similarly, overexpression of WNT10A has been shown to induce a poor prognosis^42^, and is known to interact with FZD5^43^. Finally, there is also evidence of the secretion of SECTM1 by dendritic cells to attract monocytes to the TME via binding to CD7, promoting their differentiation into macrophages^44^. Taken together, pro- or anti-tumor immune infiltration through macrophage communication with dendritic cells is more likely to occur in patients from non-desert immune phenotypes^31^.

### Network features as indicators for response to ICBs

As the graph features derived by RaCInG provided mechanistic understanding in terms of cell-cell communication about patients’ immune phenotype, we were interested in extending the analysis into investigating patients’ response to anti-PD1 immunotherapy^45–47^ (**Methods; Supplementary Tables 2**).

First, we analyzed two melanoma datasets (Gide-Aulander cohorts^45,46^) with known ICB response and RNAseq data from samples collected before (n = 51) and on (n = 26) treatment. We computed the average (theoretical) kernel-values (**Methods**) for the responder and non-responder patients and used it as a measure of direct communication between cell-types in the TME (**Fig. 7A**).

**Fig. 7:**
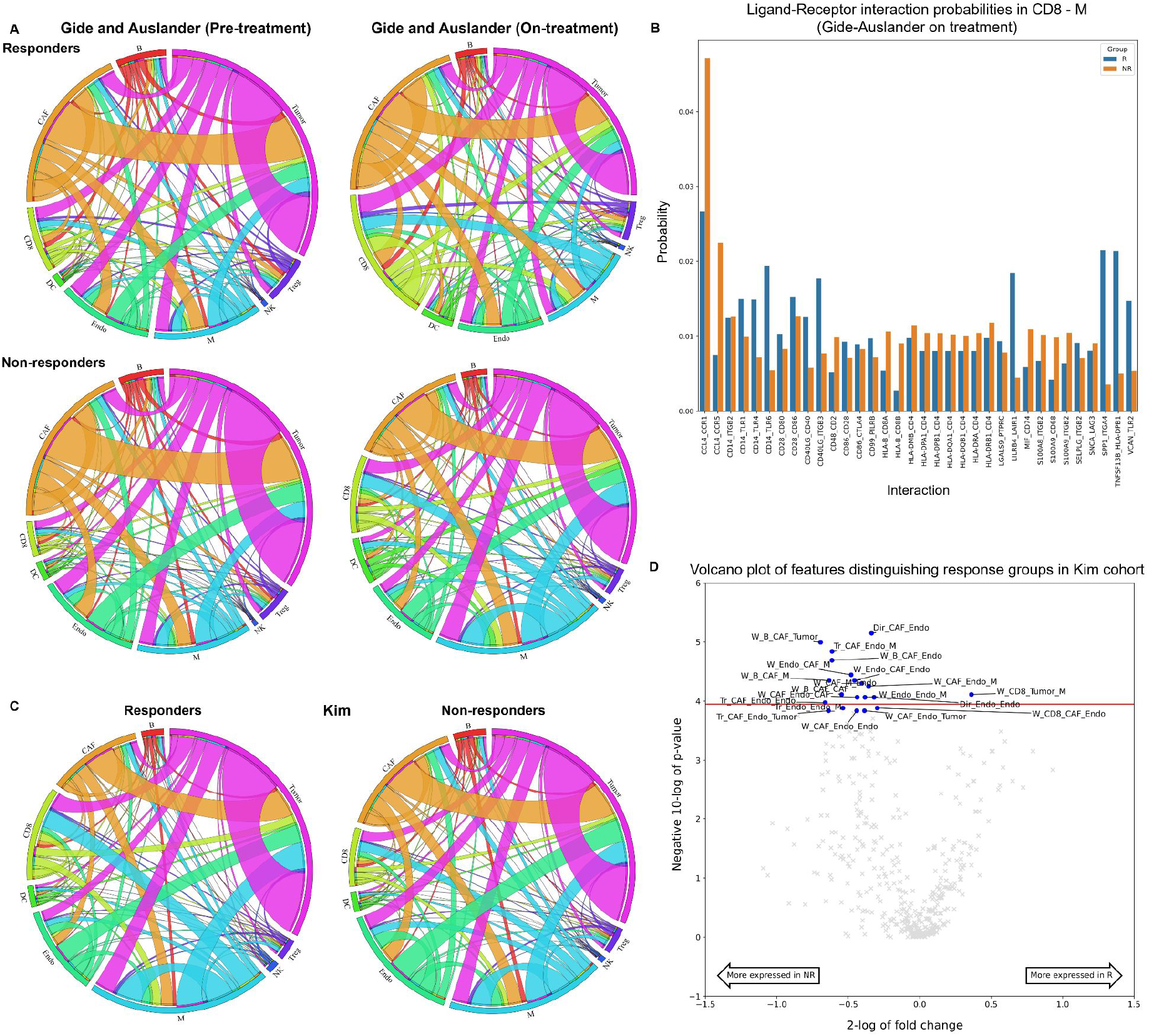
Comparison of responders and non-responders to immunotherapy. (A) Circos plot of average kernel values in responder and non-responder groups of the Gide-Auslander cohorts. The size of each ribbon indicates the fraction of total communication each cell-type is part of. The thickness of the lines in between two cell-types indicates how much these cell-types communicate. Circos plots were produced using the online tool “circos”^50^. (B) Protein communication scores between CD8+ T-cells and macrophages in the Gide-Auslander cohorts on treatment. (C) Circos plot of average kernel values in responder and non-responder groups of the Kim cohort. (D) Comparison of responders and non-responders in the Kim cohor

For the responding patients in the Gide-Auslander cohorts we observed a large increase of CD8+ T-cell communication during immunotherapy (sum of the kernel values 1.4576 vs 2.8788 for before and on-treatment samples, respectively). When we zoomed into the types of communication this cell-type was involved in, we observed specifically that both the amount of communication from tumor to CD8+ T cells (kernel value 0.2930 before treatment vs 0.4510 on treatment) and from macrophages to T-cells (0.1558 vs 0.3170) doubled. Moreover, the amount of communication in between T-cells almost quadrupled (0.1159 vs 0.4002). Overall, this suggests an increased activation of CD8+ T cells enabled by ICB therapy.

These results are in agreement with the increased CD8+ T cell communication in the IE subtype (associated with ICB response^31^) with respect to the F subtype (associated with lack of ICB response^31^) that we previously observed in the pan-cancer analysis (**Fig. 5**). For ICB to be effective, T-cell activation is important^31,48^. This means that T-cells communicate more with tumor cells (for direct killing and additive cytotoxicity^49^) or with each other (for self-activation), explaining why these two fingerprints increased in responders upon ICBs treatment.

Interestingly, when we subsequently compared non-responders before and on treatment in the same cohort, we still saw an increase in T-cell communication (the sum of all kernel values involving T-cells is 0.8212 before treatment versus 1.7490 on treatment), but not in tumor communication (5.4529 versus 5.2310). We also noted that macrophage communication to T-cells tripled (0.0924 versus 0.3398). Taken together with increased T-cell activity, this hinted at non-responders having a high activity of pro-tumor macrophages after treatment, which could not solely be explained by increased pro-tumor macrophage abundance (**Supplementary Fig. 4**). The pro-tumor macrophages possibly suppress the anti-tumor activity of CD8+ T-cells and cause resistance to ICB treatment.

To better understand the dual role that macrophages to CD8+ T cells communication have in responders and in non-responders we compared the ligand-receptor interactions driving this cell-cell interaction in the on-treatment samples between these two patient groups (**Fig. 7B**). Here, we observed that non-responders had a higher expression of the macrophage inflammatory protein 1β (CCL4), which is linked to suppression of CD8+ T-cells and recruitment of pro-tumor macrophages^51^. Moreover, we also saw increased activity of the S100 calcium-binding protein A8 and A9 (S100A8/-A9), which is a biomarker of tumor progression also in response to ICB therapy in melanoma patients, in agreement with their appearance in non-responders^52^. Additionally, non-responders showed higher expression of major histocompatibility complex class I-B (HLA-B) binding to CD8(A and B) T cell receptors, which is linked to downregulation of CD8+ T-cell activity^53^ and T-cell exhaustion^54^ mediated by pro-tumor macrophages. Interestingly, major histocompatibility complex class II (HLA-DM, -DP, -DQ, -DR) interaction with cluster for differentiation 4 (CD4), which is normally related to antigen presentation, is also slightly higher in non-responders. This could be linked to an aberrant expression of HLA class II molecules that have been linked in melanoma to recruitment of dampened CD8+ T-cells^55^. Finally, some other minor proteins that are more expressed in non-responders are also linked to pro-tumor macrophage activity (e.g., macrophage migration inhibitory factor; MIF^56^). Overall, these results suggest that macrophages to CD8+ T cells regulation in non-responders is associated with pro-tumor macrophages which provide alternative ways to inhibit immune response and therefore resist anti-PD1 treatment.

Next, we analyzed responder and non-responder gastric cancer patients treated with anti-PD1 (Kim cohort, n = 45; **Fig. 7C**)^47^. We observed that features involving CAFs and endothelial cells were indicative of non-response (e.g. *Dir_CAF_Endo* in **Fig. 7D**; p-value < 0.0001 and 2-log fold-change -0.34) while the *W_CD8_Tumor_M* feature was indicative of response (p-value < 0.0001 and fold-change 0.36). Communication of CAFs and endothelial cells could be expected for the non-response group, given their known association with angiogenesis and a pro-tumor microenvironment^57^. The appearance of CD8+ T-cell communication with tumor cells was in line with the general behavior for responders we observed when analyzing the melanoma cohorts.

Since macrophages made an appearance in both the non-responder and responder groups, we further looked at direct cell-cell communications (**Fig. 7C**). We observed that macrophages communicate more with CD8+ T-cells in the responder (kernel of 0.4483) than in the non-responder group (kernel of 0.2462). This is in agreement with previous observations that, in gastric cancer, interactions between CD8+ T-cells and macrophages create an immune inflamed TME that is associated with better prognosis and survival^58^. Additionally, in the non-responder group the protein interaction profile showed that the macrophages exhibited a pro-tumor phenotype (**Supplementary Fig. 5**).

Finally, we observed that B-cells tend to preferentially appear in features associated with non-response (**Fig. 7D**) and that they are in general more active in non-responders (sum of kernel values involving B-cells is 0.5221 in responders versus 0.7593 in non-responders; **Fig. 7C**). A possible explanation to this behavior is the formation of regulatory B-cells. This phenotype of B-cells plays a role in tumor progression and immune system suppression in gastric cancer^59^. To test this hypothesis, we compared the ligand-receptor interactions for B-cell to CD8+ T-cell communication between responders and non-responders (**Supplementary Fig. 6**). This showed that especially the lymphocyte-specific protein tyrosine kinase (Lck) was more active in the non-response group. This protein is known to hinder T-cell activation^60^, providing a pathway exploited by B-cells other than PD-L1 to allow tumor cells to evade the immune system, hinting at the regulatory B-cell phenotype^59^.

## Discussion

RaCInG is a new computational tool to construct patient-specific cell-cell interaction models based predominantly on bulk RNA-seq. This methodology leverages techniques from the mathematical field of random graphs and provides a way to build cell-cell interaction networks at the level of individual cells. Moreover, since the networks are built based on well-studied^17^ models, theoretical guarantees can be derived about the patient’s networks. Our method extends previous research efforts since it captures the unknown features in a patient’s TME through stochasticity. It assumes the input data as given but does not infer a deterministic network based on this data. Rather, it builds a network ensemble of admissible networks which adhere to the provided input data and searches for features that remain (statistically) consistent over the entire ensemble. Moreover, RaCInG allows us to go beyond communication of individual cell-types: it can consider features for which more than two cell-types come into play (wedges and triangles). Finally, RaCInG removes the bias introduced through cell-type quantification and places all network features on the same footing by considering the fold-change between feature values in two network settings: one with “normal” input data, and one with input data that only considers cell-type quantification.

We have applied RaCInG to study the role of cell-cell interactions in the TME. We have shown that RaCInG is able to extract network fingerprints for individual patient’s TME that correlate well with immune response, TME subtypes and response to ICB therapy. Intracellular communication regulates cellular phenotypes possibly explaining the dual role that certain cell-types can have in different contexts^27,61^. For example, using RaCInG in a pan-cancer analysis of six cancer types from the TCGA, we have observed that macrophages can be associated with an anti-tumor or a pro-tumor TME subtype depending if they preferably interact with CD8+ T-cells or with CAFs, respectively. In agreement with this observation, macrophages interaction with CD8+ T-cells is also positively associated with better response to ICB therapy in gastric cancer.

RaCInG allows us to dive deeper into which ligand-receptor pairs characterize specific cell-cell interactions, providing a better understanding of their potential role in regulating immune cell phenotypes. In this way we can link network fingerprints with their corresponding proteomic landscape. For example, we used this approach to look into ligand-receptor interactions driving communication between macrophages and CD8+ T cells in melanoma patients treated with ICB therapy. In this way we identified a potential role of pro-tumor macrophages in downregulating immune response in non-responders, which could justify their lack of response to anti-PD1 therapy. If this is the case, combining anti-PD1 therapy with macrophage-targeting treatment could provide a better treatment strategy.

It is worth noticing that more “complex” features (i.e., wedges and triangles, involving triplets of cell-types) are often better associated with tumor characterization than the “simpler” direct communication features between pairs of cell-types. Although cellular communications involving more than two cell-types are more difficult to interpret, this observation highlights the importance of studying cell-cell communication networks rather than focusing only on direct cell-cell interactions pairs.

In the TCGA case study we are not using the model to its full potential: we are extracting undirected features from the graph while we are constructing models that consider directionality. This choice was made based on the observation that directed features involving the same cell-types often had similar relevance, e.g., in the pan-cancer analysis comparing different TME subtypes. A possible explanation for this in the context of the TME, is the observation that ligand-receptor interactions are not a one-way street. When a ligand-receptor interaction occurs, often this elicits a reaction in both the ligand cell-type, and the receptor cell-type^10^, partially masking the directionality of the interaction.

RaCInG does not perform well with very rare cell-types like the NK cells in our case study, possibly because they cannot be accurately estimated with current deconvolution methods. In practice, this meant that NK cells might not appear in any of the networks for some patients, rendering features including NK cells unstable. To overcome this issue, we did not include NK cells in our statistical analysis for the features computed with the monte-carlo method.

For direct communication we overcame this issue by resorting to the kernel method. Kernels are derived from random graph theory and capture the limiting behavior of the networks when the amount of cells tends to infinity. Since RaCInG’s kernel is determined by a relatively simple equation, this allows us to derive feature values much quicker and more precisely than in the monte-carlo method. However, we can only apply the kernel method in cases where the exact feature values are theoretically known. When no theoretical results exist, one needs to rely on the monte-carlo method.

Although we have shown the potential of RaCInG when applied to some case studies, we provide RaCInG as a flexible and modular tool that can be adapted to different research needs. For example, we currently implemented the extraction of three types of network fingerprints (direct communication features, wedges, and triangles), however other graph fingerprints (e.g, the size of the giant strongly connected component, or more intricate graphlets like stars) can be integrated in the pipeline. RaCInG can also be adapted to using different input data, for example when cell quantification in a sample is directly measured (e.g., by flow cytometry). This information can be provided as input to RaCInG without having to rely on deconvolution methods. Similarly, when more context specific information on the expression of ligands and receptors are available for individual cell-types, this information can be directly used to compute the LR distribution matrix. Random graph methods could be expanded in the future to include geometry^20^ to leverage the increasing availability of spatial data (e.g. spatial transcriptomics^62^ or immunohistochemistry^63^).

To conclude, we envision that RaCInG will be a useful tool to study how cell-cell communication characterizes the observed tissue phenotypes in different contexts. This can extend to the investigation of intercellular communication in different physiological (e.g. cell development^64–67^ or tissue homeostasis^68^) and pathological contexts^69^.

## Methods

### Cancer specific data acquisition and transformation

In the context of modeling the TME, RaCInG requires different types of biological information. We first annotated which ligand-receptors are specific for the different cell-types of interest by leveraging curated literature resources^23^ and cell-type specific RNA-seq data^24^. And then, we used bulk RNA-seq data to quantify cell type fractions and ligand-receptor bindings for each individual patient.

To better characterize the cell-cell communication network produced by RaCInG, we gathered information about the TME subtype of patients (from literature) and their anti-cancer immune response (inferred from bulk RNA-seq).

### Bulk RNA-sequencing data

#### The Cancer Genome Atlas (TCGA)

Gene expression data for six solid tumors: BLCA, CRC, NSCLC, KIRC, SKCM and STAD were downloaded via the Firehose tool from the BROAD Institute (https://gdac.broadinstitute.org), released January 28, 2016. We selected primary tumor or metastatic (only in the case of melanoma) samples, resulting in a total of 3213 patients.

We extracted the gene expression data from “illuminahiseq_rnaseqv2-RSEM_genes” files. From these data, we used “raw_count” values as counts, and we calculated transcripts per million (TPM) from “scaled_estimate” values multiplied by 1,000,000. We first removed those genes with a non-valid HGNC symbol and then we averaged the expression of those genes with identical HGNC symbols.

#### Datasets of patients treated with immunotherapy

Gene expression data for melanoma (Gide^45^ and Auslander^46^ cohort) and gastric cancer (Kim^47^ cohort) was available from published datasets of patients treated with anti-PD1 therapy, which also include information about patients’ best overall response (**Supplementary Table 3** for more details and accession numbers).

For each cohort, we downloaded FASTQ files of RNA-seq reads from the Sequence Read Archive (SRA, https://www.ncbi.nlm.nih.gov/sra/). We used quanTIseq to process the data^70^. First, Trimmomatic ^71^ is used to remove adapter sequences and read ends with Phred quality scores lower than 20, discard reads shorter than 36 bp, and trim long reads to a maximum length of 50 bp (quanTIseq pre-processing module). Then, Kallisto^72^ is applied on the preprocessed RNA-seq reads to generate gene counts and TPM using the “hg19_M_rCRS” human reference (quanTIseq gene-expression quantification module).

### TME subtypes

We used a previously defined classification of the TME to assign patients into different subtypes: Immune-Enriched Fibrotic (IE/F), Immune-Enriched Non-Fibrotic, Fibrotic (F) and Desert (D)^31^. The TME subtype associated with each patient was provided by the original work for TCGA datasets as well as for Gide-Auslander cohorts.

### Transformation of RNA-seq into RaCInG input data

#### Quantification of individual cell-type abundance

We used in silico deconvolution^73^ to estimate cell fractions from bulk-tumor RNA-seq data. In order to obtain robust cell fraction estimates, we used a consensus approach based on six deconvolution methods accessible through the immunedeconv^74^ R package v2.1.0: quanTIseq^70^, EPIC^75^, ConsensusTME^76^, xCell^77^, TIMER^78^, and MCP-counter^79^. quanTIseq and EPIC were selected for their capability of estimating cell fractions referred to the overall composition of the tumor sample (not possible for the other methods), whereas the remaining methods were used to confirm and/or refine the estimates as explained in the following. quanTIseq was used to estimate cell fractions for CD8+ T cells, B cells, Tregs, M1 and M2 macrophages, which showed high correlation with the other deconvolution methods (**Supplementary Fig. 7**). Since M1 and M2 signatures do not recapitulate their diversity in the tumor and given the limited availability of methods to derive a consensus we decided to sum them and consider macrophages as a unique cell type. EPIC was used to estimate CAFs (absent in quanTIseq signature), NK cells (low consensus agreement for quanTIseq), and tumor cells (high agreement with quanTIseq estimates, but more accurate as they do not include endothelial and epithelial cells), and normal cells (endothelial cells). Treg and NK cell fractions that were given a null score by xCell, were set to zero. Given the low agreement of EPIC and quanTIseq on DC fractions compared to other methods, we used a three-step consensus approach: 1) we scaled in the 0-1 range DC scores obtained with xCell, MCP-counter, and TIMER; 2) we took their median; and 3) we rescale it to span the range of values covered by quanTIseq, after correction of absent cells according to xCell. Finally, cell fractions in each sample were rescaled to sum up to 1.

#### Cell-type compatibility of ligands and receptors

Using the LIANA^80^ R package v0.1.10 and the OmnipathR R package v3.7.0, we retrieved a customized set of intercellular interactions from OmniPath^23^, which consisted of interactions curated in the context of cell-cell communication available from six resources: CellphoneDB^81^, CellChat^82^, ICELLNET^83^, connectomeDB2020^84^, CellTalkDB^85^ and Cellinker^86^. Then, we filtered for direct cell-cell communication interactions by excluding proteins related to the extracellular matrix. Additionally, protein complexes were splitted into individual subunits. This resulted in a total of 3081 LR interactions.

From the database of Ramilowski et al.^24^, the gene expression of 144 human cell-types based on cap analysis of gene expression (CAGE) from the FANTOM5 project is available. We kept only the cell-types for which we could quantify their abundance based on deconvolution methods. The agreement was not perfect and certain “deconvolution” cell-types matched more than one “ramilowski” cell type, thus we aggregated them by averaging their expression because they showed high correlation between the expression of their ligands and receptors. We additionally included a pan-cancer cell type derived by using data from the Cancer Cell Line Encyclopedia (CCLE)^87^ as described in our previous study^26^. Based on gene expression data of 583 cell lines (from 18 solid cancer types), the median expression of each gene was considered as the gene expression of the pan-cancer cell type.

Ligands and receptors were first selected based on their expression (≥10 TPM threshold) in at least one of the 10 cell-types considered, and then based on the presence of the corresponding ligand or receptor pair in the network. The 10 TPM threshold was initially used in the Ramilowski paper for the CAGE data, and it was based on known expression data from B-cells. We have previously described that this cutoff value was suitable for the CCLE RNA-seq data^26^.

The compatibility of ligand and receptors was specific for each cell type, comprising a total of 971 LR pairs.

#### Quantification of ligand-receptor pair activation

Patient-specific LR pair weights were defined as the minimum of the log2(TPM+1) expression of the ligand and the receptor, hypothesizing that the expression of the gene at the lower level limits the LR binding affinity.

#### Computation of immune response score

We used our “easier” R/Bioconductor package^26,88^ to compute a score of immune response based on the median of the z-score values of 10 published transcriptomics signatures of the immune response. All these signatures were calculated according to the methodology reported by the original studies.

### Random graph generation (monte-carlo simulation)

The process in which RaCInG created graphs and extracted features is independent of the application domain. Four different facets are important in this pipeline:

1. Generation of nodes and arcs based on input data.
2. Assignment of arcs to node-pairs.
3. Feature extraction.
4. Normalization.

### Generating nodes and arcs

An overview of the variables and distributions used for the random graph model is presented in **Table 2**. These variables correspond to (elements of) the input matrices in **Fig. 1A**.

**Table 2.** Symbols used to describe the random graph model.

| Symbol | Type | Interpretation | Notes |
| --- | --- | --- | --- |
| $N$ | Number | Number of cells in one network instance. | |
| $\lambda$ | Number | Average number of interactions per cell. | |
| $Q$ | Probability distribution | Probability of cells having a given type, i.e. the cell-type quantification in the C-matrix of **Fig. 1A**. | $q_k$ is the probability of one cell having type $k$. |
| $P$ | Probability distribution | Probability of an interaction consisting of a given ligand and receptor, i.e. the ligand-receptor quantification in the LR-matrix of **Fig. 1A**. | $p_{ij}$ is the probability of one interaction consisting of ligand $i$ and receptor $j$. |
| $L$ | Matrix | Compatibility of specific cell-types with specific ligands. | $L(k, i)$ is the indicator that cell-type $k$ can secrete ligand $i$. |
| $R$ | Matrix | Compatibility of specific cell-types with specific receptors. | $R(k, i)$ is the indicator that cell-type $k$ can secrete receptor $j$. |

To create the nodes for one instance of the network, RaCInG creates a list of length *N* with independent realizations from *Q*. In this list entry *l* corresponds to the cell-type of node *l*. Similarly, to create the (unpaired) arcs for one instance of the network, RaCInG creates a list of length *λN* (rounded down) with independent realizations from *P*. Here, entry *l* of the list corresponds to a tuple that encodes both the ligand and the receptor of interaction *l*.

### Pairing nodes and arcs

To pair nodes and arcs, RaCInG iterates over the list of interactions in the following way:

1. It reads the type of the interaction’s ligand. Suppose it had type *i*.
2. It highlights all nodes that have a type *k* such that *L*(*k, i*) = 1.
3. It chooses one of these nodes uniformly at random with replacement.
4. It reads the type of the interaction’s receptor. Suppose it had type *j*.
5. It highlights all nodes that have a type *k* such that *R*(*k, j*) = 1.
6. Independently of the previous choice, it chooses one of these nodes uniformly at random with replacement.

After this procedure is executed for all interaction pairs, we obtain a complete network. To generate an ensemble of networks for one patient, RaCInG repeats the node/interaction procedure and pairing procedure a predetermined number of runs. Each run is generated independently from the previous runs.

### Feature extraction

#### Wedges and triangles (monte-carlo method)

For wedges and triangles the feature extraction is based on a network’s adjacency matrix *A*. In this matrix the entry α_í;_· indicates the number of arcs from node *i*. to node *j*. For each network, RaCInG outputs a list of paired arcs, which is transformed into an adjacency matrix. Features are then extracted from this matrix.

For example, for the wedges this is done by iterating over all rows in the matrix, recording the neighbors a given vertex connects to (together with the multiplicity of the connection) and then recording these neighbors’ subsequent neighbors. This yields a list of triplets of vertices that form wedges. The types of these wedges can subsequently be extracted and tallied for each combination of cell-types. Triangles counts are computed in a similar way.

Once this procedure is executed for each individual network in the ensemble, the average is computed over all the tallies. This provides the value of one feature for a given patient. The standard deviation is also recorded as a check to ensure the average expression value concentrates around the actual measured feature values from each network.

#### Direct communication (kernel method)

For direct communication values an asymptotic count is implemented based on the law of total probability and the law of large numbers^89^. To derive this count, we first note that the expected fraction of cells a fixed ligand *i*. can connect to, is given by ∑*_s_ q_s_L*(*s, i*). Similarly, the expected fraction of cells a fixed receptor *j* can connect to is given by ∑*_r_ q_r_R*(*r, j*). Together, ∑*_sr_ q_s_L*(*s, i*)*q_r_R*(*r, j*) is the fraction of cells an LR-pair (*i, j*) can connect to.

An arc from cell-type *k* to *l* is only formed if cells of these types are chosen in the arc assignment step. Since the fraction of cells with these types is given by *q_k_* and *q_l_*, respectively, and since only one pair of (admissible) cells is chosen uniformly at random, the probability that LR-pair (*i, j*) formed an arc from cell-type *k* to *l* is given by

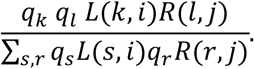

Here, multiplication with *L*(*k, i*)*R*(*l, j*) is needed since the probability can only be nonzero when the LR-pair is allowed to connect cells with type *k* and *l*. This probability is built on the assumption that LR-pair (*i, j*) is chosen to connect two cells. In reality, RaCInG can generate all possible LR-pairs to connect cells, hence it is not known a-priori. Thus, to find the a-priori probability of an arc being formed from cell-type *k* to *l*, a weighted sum needs to be taken over the above probability for all possible LR-interactions. The weight for each probability is given by the LR-pair’s quantification *p_ij_*. Mathematically, this means the law of total probability is applied, and it yielded the following a-priori probability *π_kl_* of generating a connection from cell-type *k* to *l*:

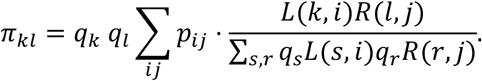

With this probability *π_kl_*, an asymptotic count can be computed. If *N_kl_* denotes the total number of arcs from cells of type *k* to *l*, then it is known due to independence of the various arc placements in RaCInG’s network generation algorithm that *N_kl_* is a binomial distribution with *λN* trials and success probability *π_kl_*. From this fact, together with the (weak) law of large numbers, we subsequently conclude that

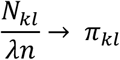

in probability. These were the theoretical feature values used for direct communication. Moreover, the expression

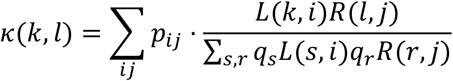

within the expression of *π_kl_* is called the kernel of RaCInG. It could be interpreted as the direct communication feature with the explicit cell-type quantification bias (the product *q_k_ q_l_*) removed.

#### From directed to undirected features

Features from both the monte-carlo and kernel method are directed. For the TCGA case study it was decided to use undirected features instead of directed features. To compute these from the directed features, all directed counts with the same cell-types were accumulated. For example, in the case of direct communication the undirected feature Dir_A_B was obtained by computing *π_AB_* + *π_BA_*. A visual overview of all directed features to accumulate to get the corresponding undirected feature is presented in **Supplementary Fig. 8**.

### Normalization

To normalize, the pipelines for network generation and feature extraction were executed again, but this time in a setting where the distribution *P* was made uniform over its support. Hence, if one sets

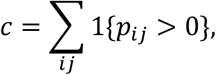

where 1{·} indicates the indicator function, then in the uniform runs a new probability distribution 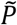 was used for the ligand-receptor interactions. In this distribution, the probability of an interaction between ligand *i*. and receptor *j* occurring was given by

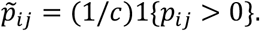

All other parameters were kept the same as in the previous “standard” runs. Finally, if *f_st_* is the (average) feature value in the “standard” run and f_unif_ is the same (average) feature value in the uniform run, then the normalized feature value was given by the fold change between these two runs, i.e.

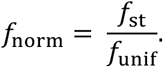

One can identify *f_norm_* as the number of times a feature would appear more often in the networks generated with the actual input data compared to the networks generated with input data that disregarded the LR-quantification. A big advantage of this normalization procedure is its ability to place all feature values on the same footing regardless of the method they were computed by. No matter if *f_norm_* is computed through the monte-carlo method or the kernel method, its interpretation and value range stay the same.

### Analyzing the extracted features

#### Statistical analysis methods

We used the Spearman rho correlation coefficient to assess correlations between two samples. This metric was applied, since limited prior knowledge was available on the joint distribution of the two samples. Moreover, since some features extracted from the networks (especially ones with small cell-type or ligand-receptor quantifications) were prone to producing outliers, a metric is used that is more robust against these outliers. To calculate it the scipy.stats.spearmanr function in Python was used based on the 1.9.2 version of the Scipy package.

The two-sided Wilcoxon rank sum test at significance level *α* = 0.05 was applied to test for differences between two groups of patients in the case studies. It was chosen for similar considerations as the spearman rho metric. To apply the test the function scipy.stats.ranksums from the 1.9.2 version of the Scipy package in Python was used. If a statistical difference between two groups was observed for a feature, the fold-change between the average feature values of the groups was used to infer how much the empirical distributions of the two groups overlap.

To correct for multiple hypothesis testing we applied Bonferroni correction by lowering the significance level for individual tests. Specifically, when we tested at significance level *α* for *n* features, the null-hypothesis was rejected whenever the test’s p-value dropped below *α/n*.

#### Bayesian computation of ligand-receptor probability for given cell-types

To compute the conditional probability that a certain LR-pair caused the formation of an interaction, given the interaction is between two given cell-types we only used the LR-distribution *P* and the compatibility matrices *L* and *R*. The unconditional probability of LR-pair (*i, j*) appearing is given by *p_ij_*. To infer its contribution to a direct interaction between cell-type *fc* and *l*, one first needs to know whether it connects these cell-types. The indicator of this event is given by *L*(*k, i*)*R*(*i, j*).

Now, since all interactions were sampled and paired independently, and uniformly at random, the conditional probability of LR-pair (*i, j*) connecting cell-types *fc* and *l* was given by the LR-pair’s relative (probabilistic) weight when compared to the weights of all LR-pairs that can connect cell-types *k* and *l*. Thus, the conditional probability that LR-pair (*i, j*) formed a connection, given that it is a connection between cell-types *k* and *l*, is given by

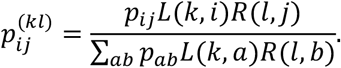

To compute the LR-probability for given cell-types over an entire group, these probabilities were taken for all patients in the group and averaged. The largest of the resulting averages were depicted in the LR-interaction bar charts.

## Data and code availability

All the datasets used are publicly available (**Supplementary Table 3**).

The code used for generating the random graphs is available at https://github.com/SysBioOncology/RaCInG. A step-by-step reproducible report (i.e., RMarkdown notebook) on how this knowledge can be extracted is made available in github. A demo that showcases RaCInG’s functionalities is also made available in github.

## Acknowledgements

The authors acknowledge the support of the Immunoengineering program of the Institute for Complex Molecular System. We would like to thank Livy Nijhuis for testing the code.

## Author contribution

PvdH and FE designed the research. MvS, OLS, PvdH and FE discussed how to use random graph models in biological context. MvS defined and implemented the mathematical formulation of the model under the supervision of PvdH. OLS analyzed the data used for the case study and transformed it into input matrices for the mathematical model under the supervision of FE. FF defined the approach for cell-type quantification using an ensemble of deconvolution algorithms. MvS, OLS, PvdH and FE contributed to the interpretation of the results. MvS and FE co-wrote the manuscript with input from all authors. All authors discussed the results and commented on the manuscript.

## Competing interests

The authors declare no competing interests.

## Notes

### Competing Interest Statement

The authors have declared no competing interest.

