## Supplementary Information for "Mathematically mapping the network of cells in the tumor microenvironment"

**Supplementary Table 1: Feature outputs from RaCInG after its application to six solid cancers from TCGA.** The first column contains patient identifiers, the last the associated cancer type, the one before the immune response score, and the one before the immune subtype identifier. All the other columns contain the feature values as computed by RaCInG for a specific feature in our test.

*SupplementaryTable1_Model_output_unknown_response_immunotherapy.csv*

**Supplementary Table 2: Feature outputs from RaCInG after its application to two cancers with known response to immunotherapy.** The first column contains patient identifiers, the last the cancer cohort, and the one before an indication whether the patient responded to immunotherapy. All the other columns contain the feature values as computed by RaCInG for a specific feature in our test.

*SupplementaryTable2_Model_output_known_response_immunotherapy.csv*

**Supplementary Table 3: Information associated with the datasets of patients treated with immunotherapy.** FFPE: Formalin-fixed paraffin-embedded; FF: Fresh-frozen; CR: Complete Responder; PR: Partial Responder; PD: Progressive Disease; SD: Stable Disease; R: Responder; NR: Non-responder.

| **Original study** | **Cancer type** | **Prior therapies** | **Biopsy** | **Samples used** | **R and NR** | **RNA-seq fastq files** |
|---|---|---|---|---|---|---|
| Auslander [1] | Melanoma (Metastasis) | Therapy naive | FF | PD-1:<br>- Pre: n=9 (R=1, NR=8)<br>- On: n=17 (R=0, NR=17) | As reported. | BioProject ID: PRJNA476140 |
| Gide [2] | Melanoma (Metastasis) | BRAFi | FFPE | PD-1:<br>- Pre: n=41 (CR=4, PR=15, PD=16, SD=6)<br>- On: n=9 (CR=0, PR=4, PD=4, SD=1) | R=CR,PR<br>NR=SD,PD | BioProject ID: PRJEB23709 |
| Kim [3] | Gastric cancer (Metastasis) | Prior failure of at least 1 line of chemotherapy (platinum) | FF | Pre: n=45 (CR=3, PR=9, PD=18, SD=15) | R=CR,PR<br>NR=SD,PD | BioProject ID: PRJEB25780 |

**Supplementary Fig 1. List of top ten features for SKCM and STAD that changed the most in average fold-change when compared to the results of the pan cancer analysis.** The direction of the triangles indicates the direction of the fold-change shift when compared to the pan-cancer analysis. Dots indicate that the fingerprint is not part of the top ten features for the given cancer type. Comparisons that do not involve the IE immune phenotype are plotted (the others are in Fig. 6).

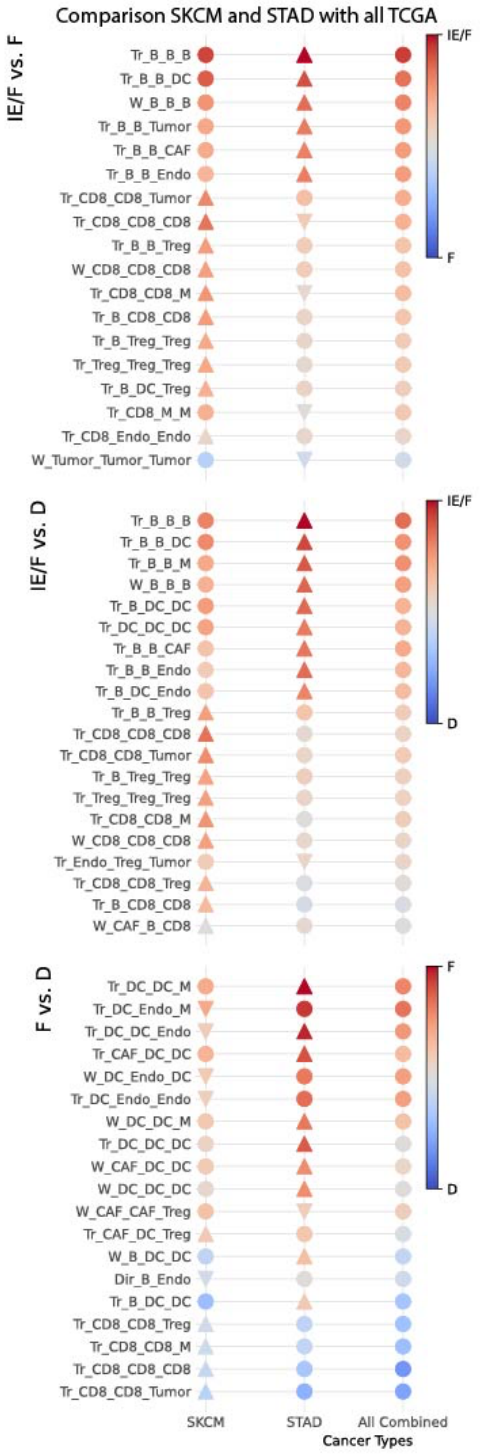

**Supplementary Fig 2. Network-based characterisation of microenvironment subtypes in SKCM.** Volcano plots showing the statistical comparison of network-based features identified by RaCInG when doing pairwise comparisons of microenvironment subtypes in SKCM. The red line indicates the $\alpha = 0.05$ significance threshold after Bonferroni correction. On the x-axis we show the fold change between the average feature values for each group, and on the y-axis the negative 10-log of the Wilcoxon rank sum test's p-value. For each plot, the twenty features with lowest p-value have been highlighted.

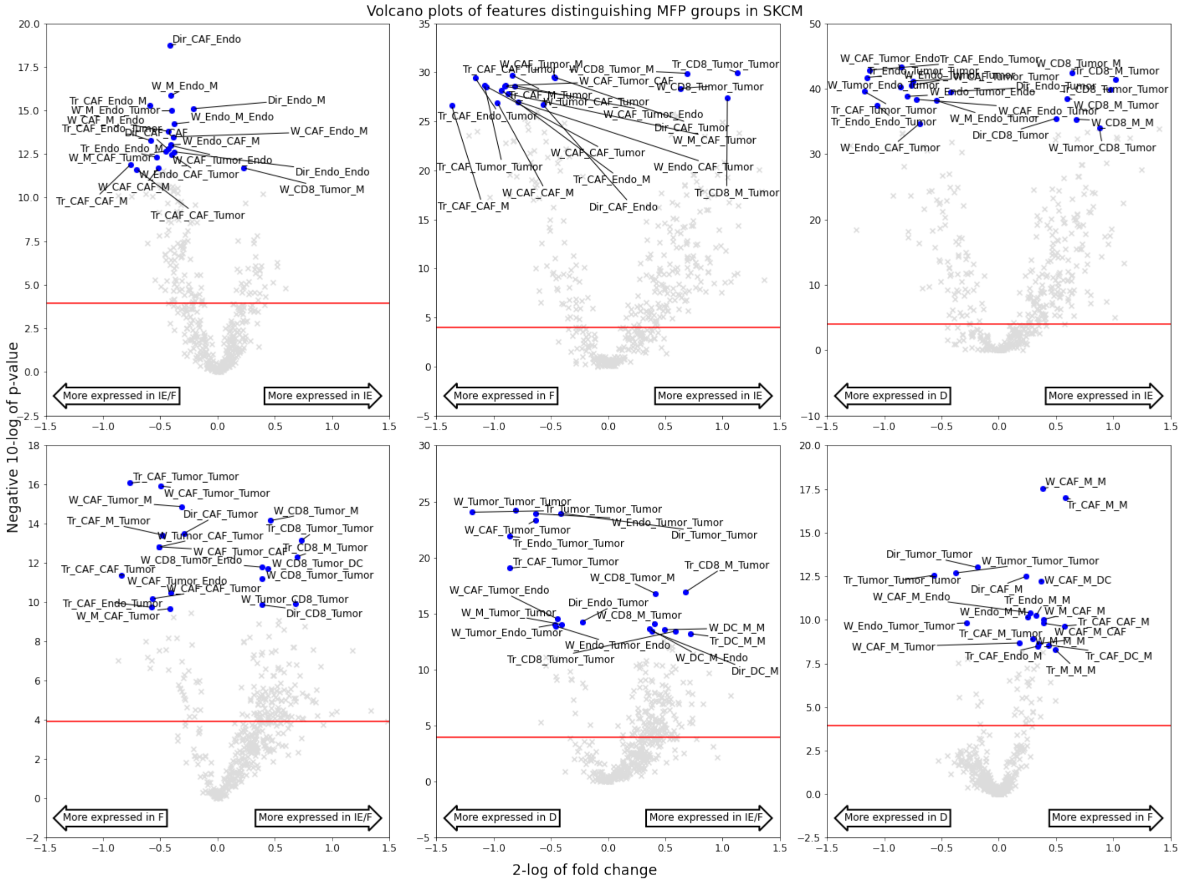

**Supplementary Fig 3. Network-based characterisation of microenvironment subtypes in STAD.** Volcano plots showing the statistical comparison of network-based features identified by RaCInG when doing pairwise comparisons of microenvironment subtypes in STAD. The red line indicates the $\alpha = 0.05$ significance threshold after Bonferroni correction. On the x-axis we show the fold change between the average feature values for each group, and on the y-axis the negative 10-log of the Wilcoxon rank sum test's p-value. For each plot, the twenty features with lowest p-value have been highlighted.

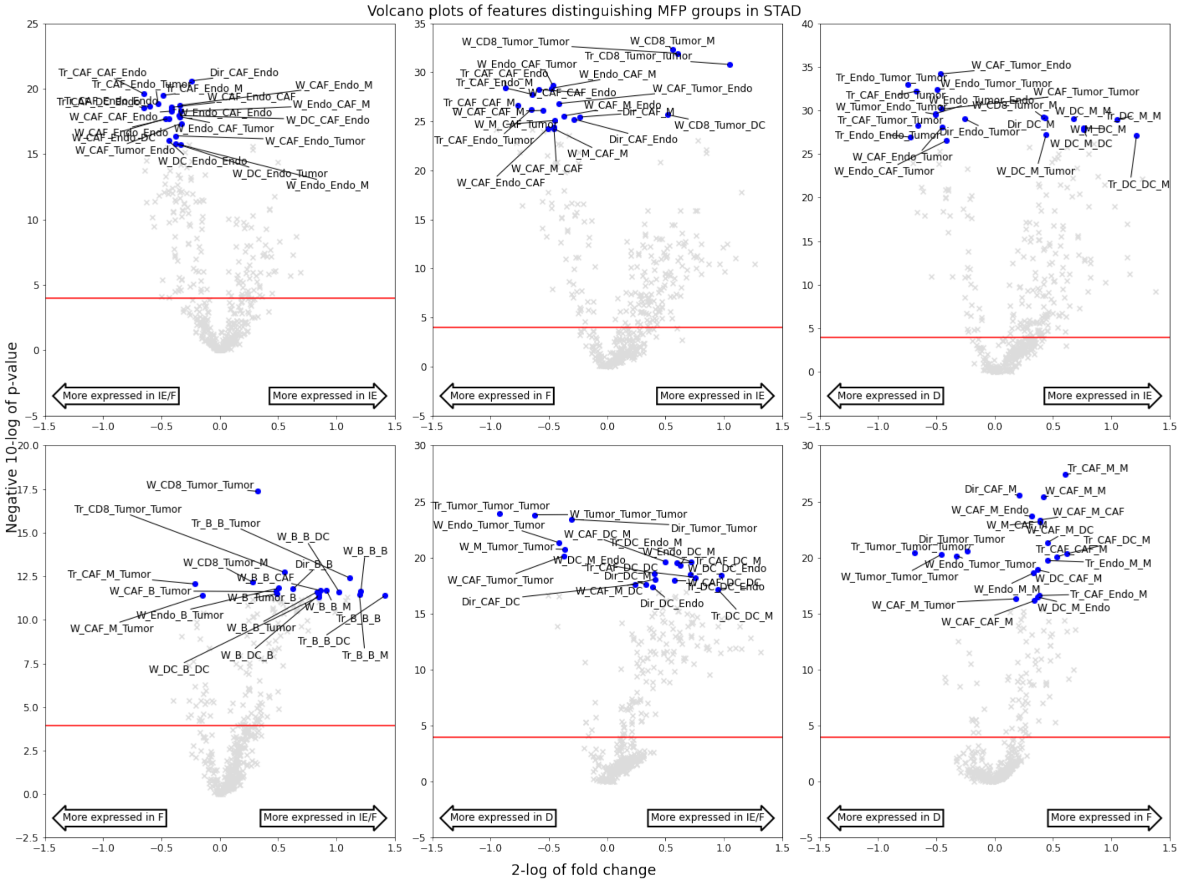

**Supplementary Fig 4. Cell-type quantification of macrophage phenotypes in patients of the Gide and Auslander cohorts before and on treatment.** Each boxplot represents the macrophage quantification of a certain phenotype in a certain response group. The boxplots have been generated for the patients before and on immunotherapy.

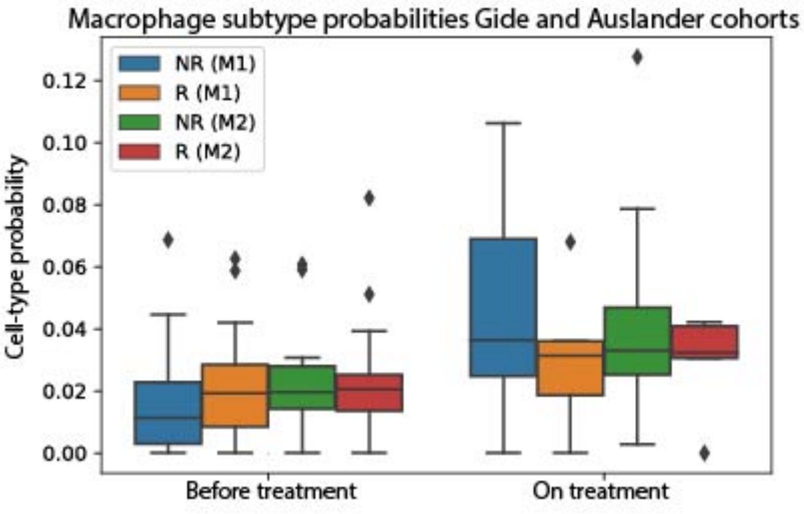

**Supplementary Fig 5. Protein communication scores between CD8+ T-cells and macrophages in the Kim cohort.** For each responder group the top twenty ligand-receptor pairs with the highest probability of appearing in the cell-cell interaction network have been presented.

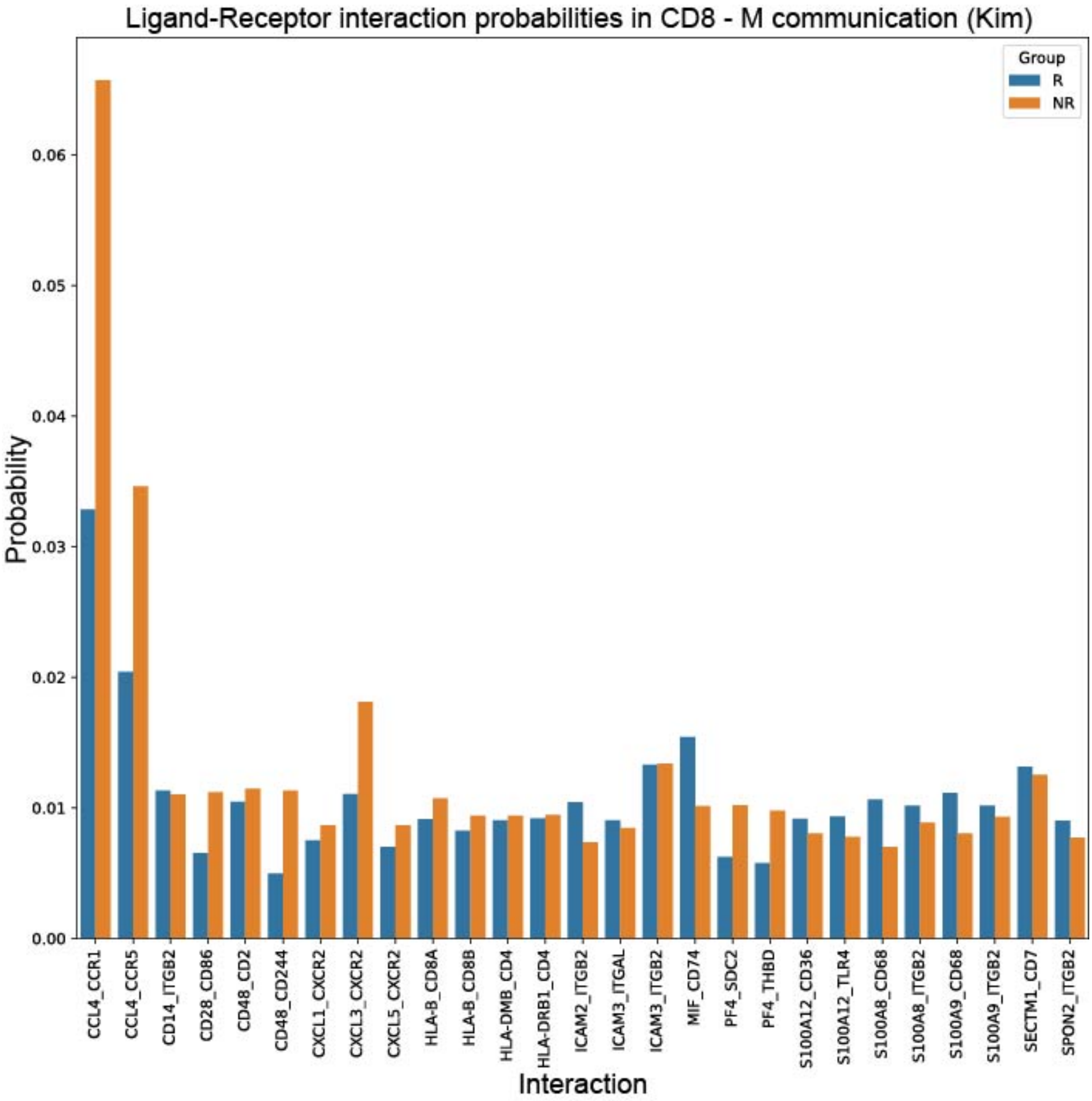

**Supplementary Fig 6. Protein communication scores between B-cells and macrophages in the Kim cohort.** For each responder group the top twenty ligand-receptor pairs with the highest probability of appearing in the cell-cell interaction network have been presented.

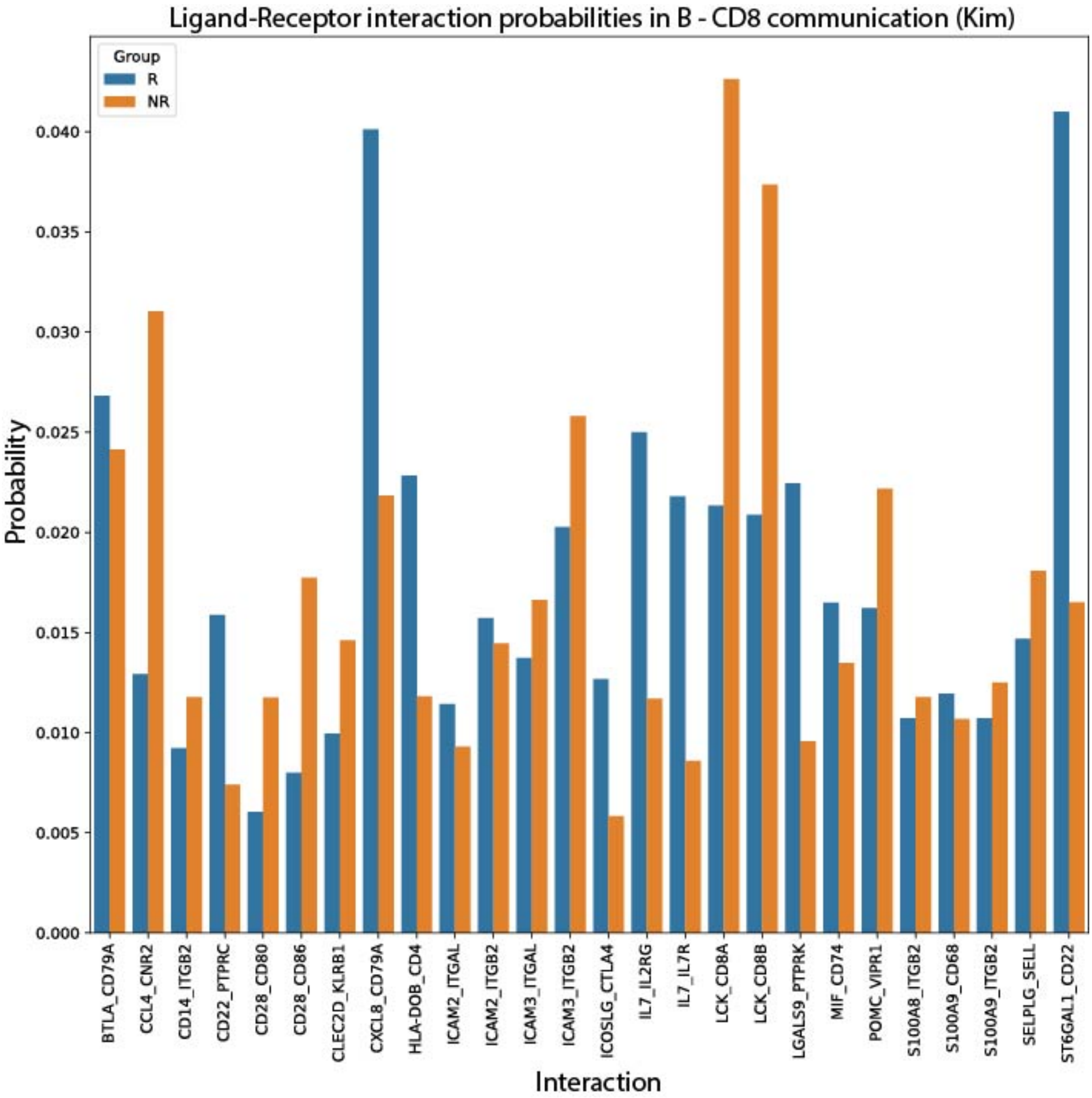

**Supplementary Fig 7. Agreement between deconvolution methods.** Mean correlation (across TCGA cancer types) between cell-type quantification computed using multiple in silico deconvolution methods.

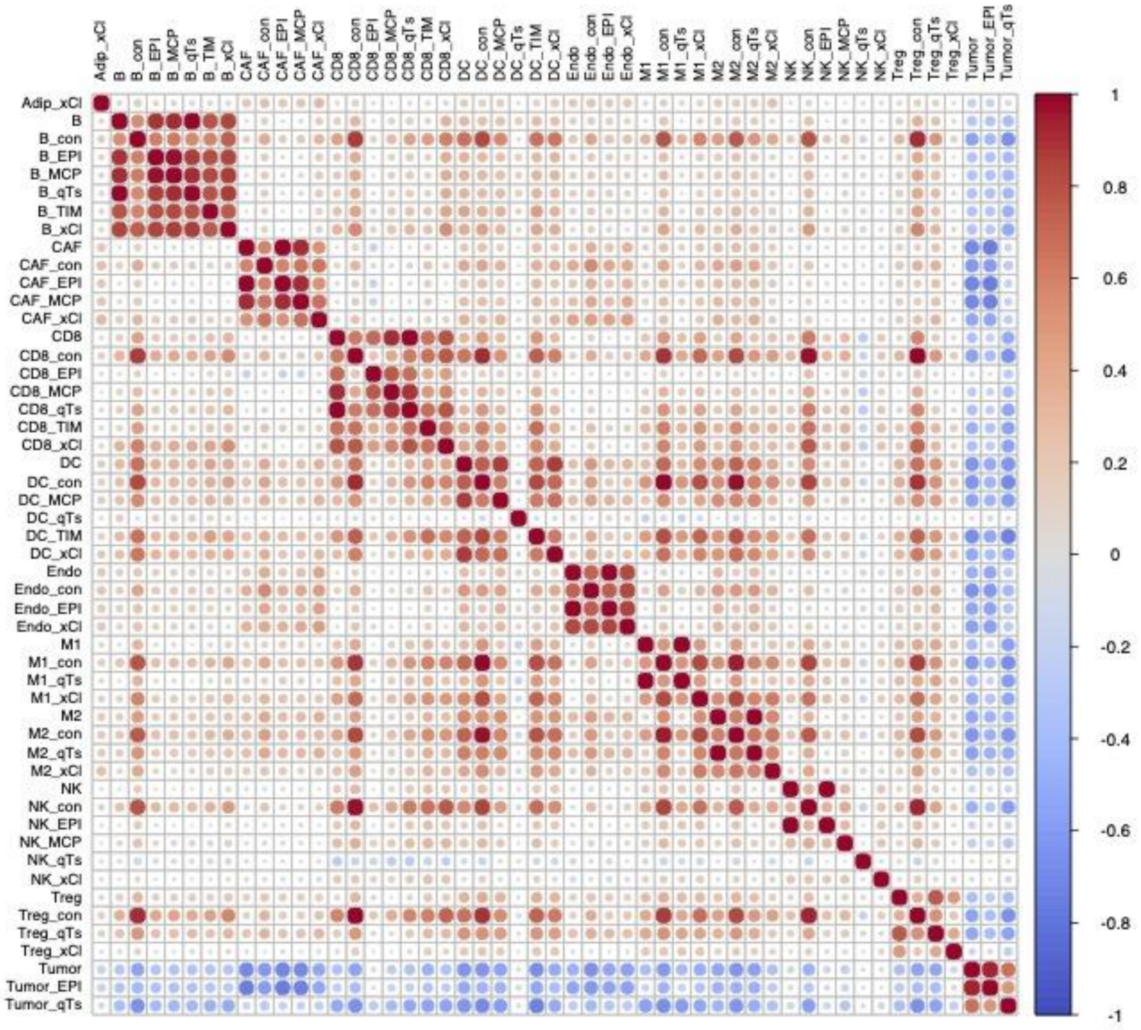

**Supplementary Fig 8. The network fingerprints that should be combined to turn directed feature values into undirected feature values.** Accumulation is done by adding the directed features values together.

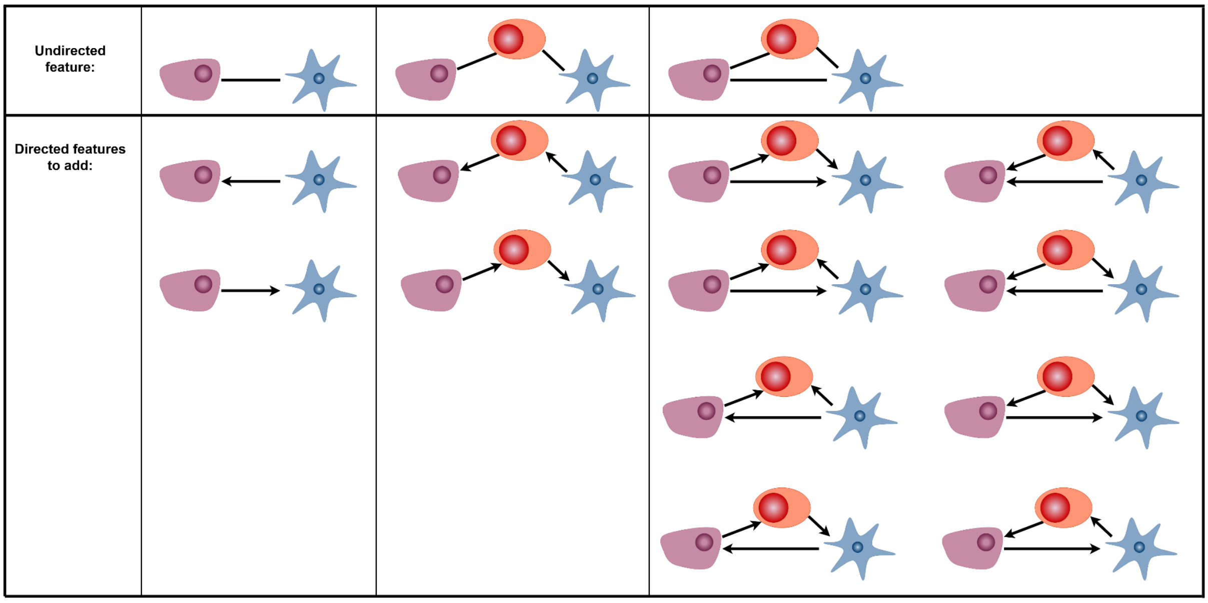
